# A robust approach to estimate relative phytoplankton cell abundance from metagenomes

**DOI:** 10.1101/2021.05.28.446125

**Authors:** Juan José Pierella Karlusich, Eric Pelletier, Lucie Zinger, Fabien Lombard, Adriana Zingone, Sébastien Colin, Josep M. Gasol, Richard G. Dorrell, Eleonora Scalco, Silvia G. Acinas, Patrick Wincker, Colomban de Vargas, Chris Bowler

## Abstract

Phytoplankton account for >45% of global primary production, and have an enormous impact on aquatic food webs and on the entire Earth System. Their members are found among prokaryotes (cyanobacteria) and multiple eukaryotic lineages containing chloroplasts. Phytoplankton communities are generally studied by PCR amplification of bacterial (16S), nuclear (18S) or chloroplastic (16S) rRNA marker genes from DNA extracted from environmental samples. However, our appreciation of phytoplankton abundance or biomass is limited by PCR-amplification biases, rRNA gene copy number variations across taxa, and the fact that rRNA genes do not provide insights into metabolic traits such as photosynthesis. In addition, rRNA marker genes fail to capture both cyanobacteria and photosynthetic eukaryotes simultaneously. Here, we targeted the photosynthetic gene *psbO* from metagenomes to circumvent these limitations: the method is PCR-free, and the gene is universally and exclusively present in photosynthetic prokaryotes and eukaryotes, mainly in one copy per genome. We applied and validated this new strategy with the *Tara* Oceans datasets, and showed improved correlations with flow cytometry and microscopy than when based on rRNA genes. Furthermore, we revealed unexpected features of the ecology of these organisms, such as the high abundance of picocyanobacterial aggregates and symbionts in the ocean, and the decrease in relative abundance of phototrophs towards the larger size classes of marine dinoflagellates. To facilitate the incorporation of *psbO* in molecular-based surveys, we compiled a curated database of >18,000 unique sequences. Overall, *psbO* appears to be a promising new gene marker for molecular-based evaluations of entire phytoplankton communities.

## Introduction

Photosynthetic plankton, or phytoplankton, consist of unicellular organisms of diverse evolutionary history and ecology. They are responsible for more than 45% of Earth’s primary production (Field, Behrenfeld, Randerson, & Falkowski, 1998), fueling aquatic food webs, microbial decomposition, and the global ocean biological carbon pump (Guidi et al., 2009). They include prokaryotes (cyanobacteria) and multiple eukaryotic lineages that acquired photosynthesis either through the primary endosymbiosis of cyanobacteria, or (and predominantly) through secondary and higher endosymbioses of eukaryotic algae (Pierella Karlusich, Ibarbalz, & Bowler, 2020). They display a broad body size spectrum, from less than 1 micron (e.g., *Prochlorococcus, Ostreococcus*) to several millimetres (e.g., *Trichodesmium* colonies, colonial green algae, and chain-forming diatoms), either due to cell size variation, aggregation or symbioses (Beardall et al., 2009). This size variability partly explains their different roles in the food web and in the biological carbon pump. For example, cyanobacteria are generally thought to be recycled within the microbial loop, whereas larger eukaryotic phytoplankton are usually considered more important in energy transfer to higher trophic levels (through grazing by small protists, zooplankton, and/or larvae (Ullah, Nagelkerken, Goldenberg, & Fordham, 2018)) and in sequestering atmospheric CO2 to the ocean interior through gravitational sinking of particles (Guidi et al., 2009).

Surveys of the structure and composition of microbial communities are typically performed by PCR amplification and sequencing of a fragment of the small subunit of the rRNA gene from an environmental sample (rRNA gene metabarcoding). The fraction of the obtained sequencing reads corresponding to a given taxon is then used as a proxy for its relative abundance. Most studies have so far focused on taxonomically informative fragments of the hypervariable regions of the 16S (prokaryote and chloroplast) or 18S (eukaryotic nuclear) rRNA genes that are by far the most represented in reference databases (Guillou et al., 2013; Pawlowski et al., 2012; Quast et al., 2013). These markers are occasionally targeted in both DNA and RNA to exclude inactive microbes and as proxies of metabolic activities (Campbell, Yu, Heidelberg, & Kirchman, 2011; Logares et al., 2012), but more recent studies have indicated severe limitations of this concept and only mRNA can be considered as an indicator of the metabolic state (Blazewicz, Barnard, Daly, & Firestone, 2013).

Although rRNA gene metabarcoding is widely used, it has multiple limitations (in addition to the error sources during DNA extraction or sequencing that also affect other molecular methods). Firstly, PCR amplification bias due to mismatches of universal primers on the target sites of certain taxa can generate differences between the observed and the genuine relative read abundances as large as 10-fold, either when using the 16S (Parada, Needham, & Fuhrman, 2016; Polz & Cavanaugh, 1998; Wear, Wilbanks, Nelson, & Carlson, 2018) or 18S rRNA gene markers (Bradley, Pinto, & Guest, 2016). Shotgun sequencing is a PCR-free alternative and consists of the detection of these marker genes in metagenomes (Liu, Lozupone, Hamady, Bushman, & Knight, 2007; Logares et al., 2014; Obiol et al., 2020).

Secondly, the copy-number of these marker genes varies greatly among species. While bacterial genomes contain between one and fifteen copies of the 16S rRNA gene (Acinas, Marcelino, Klepac-Ceraj, & Polz, 2004; Kembel, Wu, Eisen, & Green, 2012; Větrovský & Baldrian, 2013), protists can differ by >5 orders of magnitude in their 18S rRNA gene copy numbers, from 33,000 in dinoflagellates to one in small chlorophytes (de Vargas et al., 2015; Godhe et al., 2008; Mäki, Salmi, Mikkonen, Kremp, & Tiirola, 2017; Zhu, Massana, Not, Marie, & Vaulot, 2005). Due to a positive association between rRNA gene copy number and cell size, it was proposed that the rRNA gene metabarcoding reads reflect the relative biovolume proportion for a given taxon. Biovolume is a proxy of biomass, which is a relevant variable for studies of energy and matter fluxes such as food web structures and biogeochemical cycles. However, there is still little consensus for the use of rRNA gene as a biovolume estimator due to the poor correlations reported in many studies (Lamb et al., 2019; Lavrinienko, Jernfors, Koskimäki, Pirttilä, & Watts, 2021; Santoferrara, 2019; van der Loos & Nijland, 2021). Instead, there are attempts to infer relative cell abundances from rRNA gene metabarcoding by correcting the copy number variation. Although the copy number remains unknown for most microbial species, its assessment in different organisms could lead to the establishment of correction factors by assuming that the copy number is phylogenetically conserved. These approaches were applied for 16S rRNA gene in bacteria, but their accuracy is limited for taxa with no close representatives in reference phylogenies (Kembel et al., 2012; Louca, Doebeli, & Parfrey, 2018; Starke, Pylro, & Morais, 2020). In protists, this correction is even more challenging due to intraspecies variation in 18S rRNA gene copy number. For example, it varies almost 10-fold among 14 different strains of the haptophyte *Emiliana huxleyi* (Gong and Marchetti 2019). In addition, there are major difficulties for generating a comprehensive database of 18S rRNA copy numbers (Gong & Marchetti, 2019).

Finally, functional traits such as photosynthesis cannot be inferred solely from rRNA genes or other housekeeping markers, whereby their knowledge is limited to a restricted number of taxa known from experts and the literature. Indeed, while photosynthesis occurs in almost all cyanobacteria (except a few symbiotic lineages that have lost it (Nakayama et al., 2014; Thompson et al., 2012)), it is not necessarily conserved within protist taxa, such as dinoflagellates – of which only around half of known species are photosynthetic (Dorrell & Smith, 2011; Saldarriaga, Taylor, Keeling, & Cavalier-Smith, 2001) –, chrysophytes (Dorrell et al., 2019; Dorrell & Smith, 2011) and apicomplexa (Moore et al., 2008). This is an important issue because we still do not know how extended among related lineages are the independent events of chloroplast gains and losses or the extent of loss of photosynthesis with retention of the plastids. Thus, it is not possible to annotate the photosynthesis trait to those sequences whose taxonomic affiliation is, for example, “unknown dinoflagellate”.

Another disadvantage is the impossibility of making direct comparisons between cyanobacteria and eukaryotic phytoplankton with two different rRNA marker genes. This can be still attempted by targeting the plastidial and cyanobacterial versions of the 16S rRNA gene (Nicholas J. Fuller et al., 2006; N. J. Fuller et al., 2006; Kirkham et al., 2011, 2013; Lepère, Vaulot, & Scanlan, 2009; McDonald, Sarno, Scanlan, & Zingone, 2007; Shi, Lepère, Scanlan, & Vaulot, 2011). However, dinoflagellates and chromerids are not represented in these surveys because their plastidial 16S rRNA genes are extremely divergent (Green, 2011), and this approach can still capture non-photosynthetic plastids and kleptoplastids (functional plastids temporarily retained from ingested algal prey). Plastid-encoded markers directly involved in photosynthesis have also been used, such as *psbA* and *rbcL* (Man-Aharonovich et al., 2010a; Paul, Alfreider, & Wawrik, 2000; Zeidner, Preston, Delong, Massana, Post, Scanlan, & Beja, 2003). The *psbA* gene encodes the D1 protein of photosystem II and is also found in cyanophages (viruses) and the used primers target essentially the cyanobacterial and cyanophage sequences (Adriaenssens & Cowan, 2014). The *rbcL* gene encodes the large subunit of the ribulose-1,5-diphosphate carboxylase/oxygenase (RuBisCO). There are multiple *rbcL* types, even in non-photosynthetic organisms, and the gene location varies: form I is plastid encoded in plants and most photosynthetic protists (and is present in cyanobacteria) while form II is nuclear-encoded in peridinin dinoflagellates and chromerids (and is also present in proteobacteria) (Tabita, Hanson, Satagopan, Witte, & Kreel, 2008). The different *rbcL* variants thus prevent its use for covering the whole phytoplankton community.

Plastid-encoded genes (16S rRNA, *psbA*, *rbcL*) are affected by copy number variability among taxa not only at the level of gene copies (for example, four 16S rRNA gene copies in the plastid genome of the euglenophyte *Euglena gracilis* and six in the prasinophyte *Pedinomonas minor* (Decelle et al., 2015)), but also at the level of plastid genomes per plastid, and plastids per cell. The plastid number per cell varies from one or a few in most microalgal species to more than 100 in many centric diatoms (Decelle et al., 2015). In addition, this varies according to biotic interactions, e.g., the haptophyte *Phaeocystis* has two plastids in a free-living stage but increases up to 30 when present as an endosymbiont of radiolarians (Decelle et al., 2019). Photosynthetic eukaryotes typically maintain 50–100 plastid genome copies per plastid, but there is a continuous increase throughout development and during cell cycle progression (Armbrust & Virginia Armbrust, 1998; Coleman & Nerozzi, 1999; Hiramatsu, Nakamura, Misumi, Kuroiwa, & Nakamura, 2006; Koumandou & Howe, 2007; Oldenburg & Bendich, 2004). These limitations of plastid-encoded marker genes can be circumvented by the use of photosynthetic nuclear-encoded genes, which is still an unexplored approach.

In spite of the aforementioned biases, gene metabarcoding either based on rRNA genes or on alternative marker genes such as *psbA* or *rbcL* usually assume that the relative abundance of the gene sequences is an accurate measure of the relative abundance of the organisms containing those sequences. However, this assumption can lead to misleading inferences about microbial community structure and diversity, including relative abundance distributions, estimates of the abundance of different taxa, and overall measures of community diversity and similarity (Bachy, Dolan, López-García, Deschamps, & Moreira, 2013; Egge et al., 2013; Kembel et al., 2012; Mäki et al., 2017; Medinger et al., 2010; Pinto & Raskin, 2012). For example, less than 30% of the variance in true organismal abundance is explained by observed prokaryotic 16S rRNA gene abundance in some simulation analyses (Kembel et al., 2012). In addition, comparative studies between morphological and molecular approaches in environmental samples or in mock communities revealed discrepancies up to several orders of magnitude among protist taxa with regard to their relative abundances (Bachy et al., 2013; Egge et al., 2013; Mäki et al., 2017; Medinger et al., 2010; J. Pawlowski, Lejzerowicz, Apotheloz-Perret-Gentil, Visco, & Esling, 2016). Most of these studies focused on the biases generated by primers and copy-number variations, but not on uncertainties in assigning photosynthetic potential (e.g., differentiating between functionally photosynthetic and secondarily non-photosynthetic species).

We deemed it important to find more accurate alternative procedures to the most widely-used molecular approaches to make reliable estimations of species abundance, an important measure for inferring community assembly processes. We propose to target nuclear-encoded single-copy core photosynthetic genes obtained from metagenomes to circumvent these limitations: the method is PCR-free, and the genes are present in both prokaryotes and eukaryotes, in one copy per genome. We focused on the *psbO* gene, which encodes the manganese-stabilising polypeptide of the photosystem II oxygen evolving complex, and is essential for photosynthetic activity and has the additional advantage of lacking any non-photosynthetic homologs. We applied and validated this new strategy with the *Tara* Oceans datasets (Table I). We quantified the biases in taxon abundance estimates using rRNA gene markers as compared to optical approaches (flow cytometry, microscopy), and we compared these patterns with those obtained by our proposed method. We also searched for *psbO* within metatranscriptomes to analyse its potential use as a proxy of photosynthetic activity and/or biovolume (due to the higher transcript level requirements in larger cells). Besides finding a more relevant marker gene for phytoplankton, we also propose its combination with single-copy housekeeping genes (e.g., *recA* for bacteria and genes encoding ribosomal proteins in eukaryotes) to estimate the fraction of photosynthetic members in the whole community or in a given taxon. In this context, we also quantified the unknowns in the functional assignation of photosynthetic capacity based on the 18S rRNA gene. Finally, we show how the approach improves measures of microbial community diversity, structure, and composition as compared to rRNA gene metabarcoding.

**Table I:**
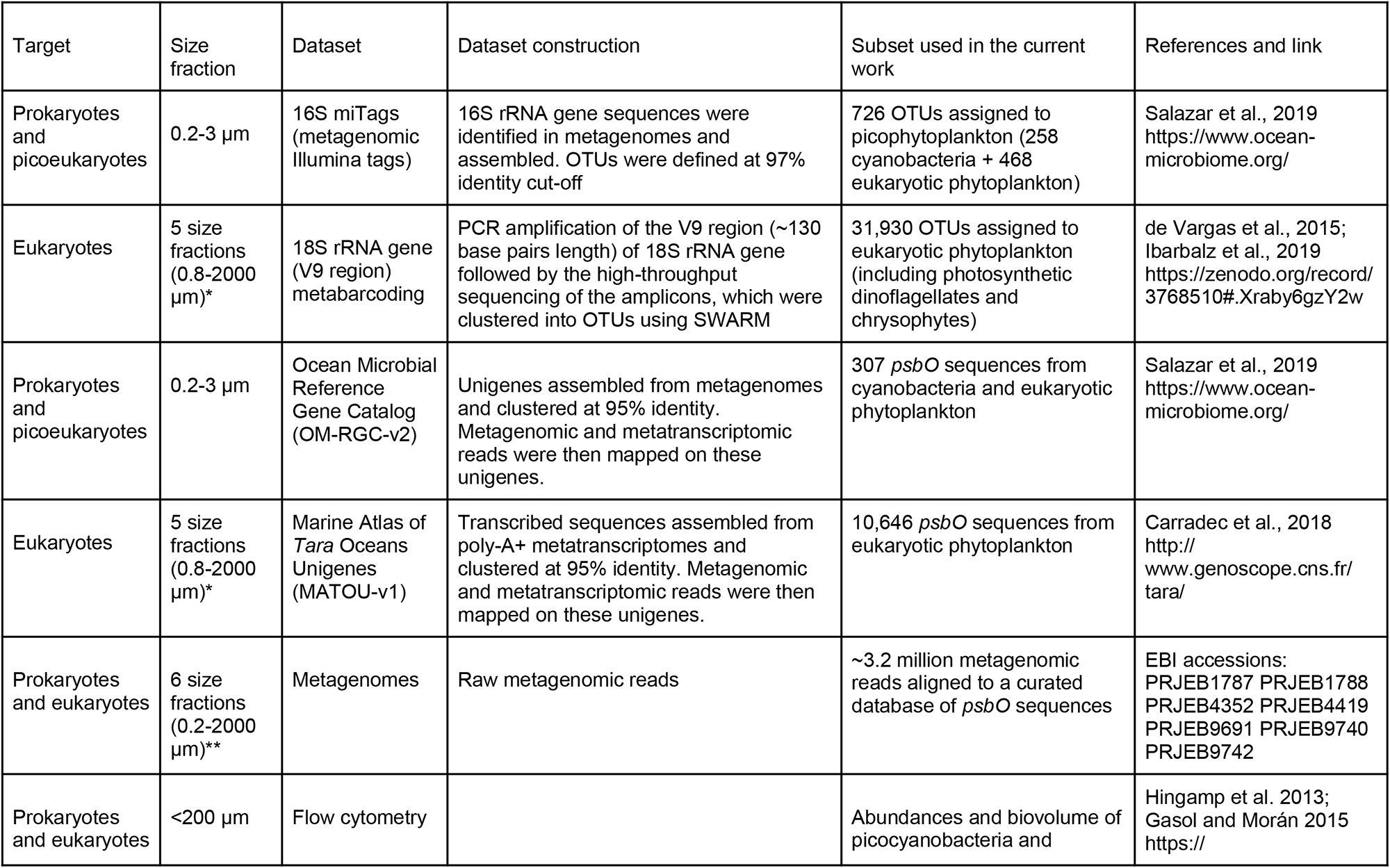
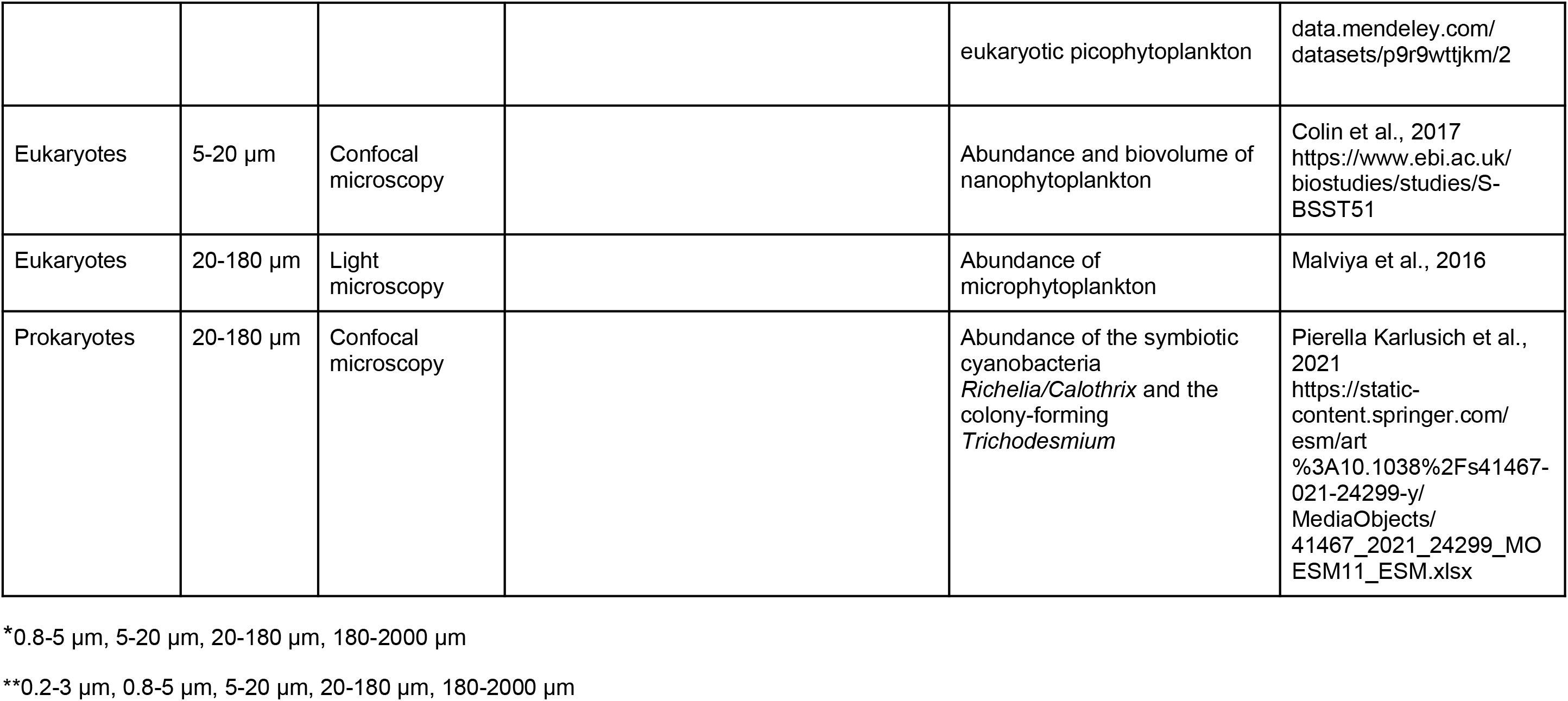
Tara Oceans datasets relevant to the current study.

## Materials and Methods

### Search for phytoplankton marker genes

To estimate cell-based relative abundances of the major marine phytoplankton groups, we searched for genes present in all photosynthetic organisms (both prokaryotes and eukaryotes) and with low copy-number variability among taxa. To fulfil the latter requirement, we first excluded the plastid-encoded genes to avoid the variations in number of chloroplasts per cell and in number of chloroplast genomes per organelle. We did this by retrieving sequences from the KEGG (M. Kanehisa, 2000) database that are assigned to the photosynthetic electron transport chain, the Calvin Cycle and chlorophyll biosynthesis to be used as queries for sequence similarity searches against >4,100 plastid genomes available at NCBI (https://www.ncbi.nlm.nih.gov/genome/organelle/). For this, BLAST version 2.2.31 (“tBLASTn” program) searches were conducted with an e-value cutoff of 1e-20 (Camacho et al., 2009). To retain only core photosynthetic genes, i.e., those present in all phototrophs, we then made an equivalent BLAST search against cyanobacterial and eukaryotic nuclear genomes from the IMG (Chen et al., 2019) and PhycoCosm (Grigoriev et al., 2021) databases and from the polyA-derived transcriptomes of the Marine Microbial Eukaryote Transcriptome Sequencing Project (MMETSP) (Keeling et al., 2014). To minimize false-negative cases, only completely sequenced genomes were considered for establishing the gene absence. This survey was also used for determining gene copy number variation.

This survey resulted in a list of five genes that are core, nuclear-encoded and present in low-copy numbers (Table II). For selecting a gene marker of phytoplankton among them, we carried out a deeper sequence analysis to detect non-photosynthetic homologs and to see if the phylogeny reflects the evolutionary history of cyanobacteria and endosymbiosis. We first performed a sequence similarity search using HMMer version 3.2.1 with gathering threshold option (http://hmmer.org/) for the corresponding Pfam domain against the translated sequenced genomes and transcriptomes from PhycoCosm and MMETSP as well as in the whole IMG database (including viruses, archaea, bacteria and non-photosynthetic eukaryotes). The Pfams used in the search were: MSP (PF01716) for PsbO, Rieske (PF00355) for PetC, PRK (PF00485) for phosphoribulokinase, UbiA (PF01040) for chlorophyll-*a* synthase, and NAD_binding_1 (PF00175) for ferredoxin:NADP^+^ reductase. CDHIT version 4.6.4 (W. Li & Godzik, 2006) was used at a 80% identity cut-off to reduce redundancy. These sequences were used for building a protein similarity network using EFI-EST tool (Zallot, Oberg, & Gerlt, 2019) and Cytoscape visualization (Shannon et al., 2003), and BlastKOALA with default parameters for functional annotation (Minoru Kanehisa, Sato, & Morishima, 2016). These analyses led us to focus on *psbO* as a gene marker for phytoplankton, for which we did a deeper analysis by building its phylogeny in the following way. Protein sequences were aligned with MAFFT version 6 using the G-INS-I strategy (Katoh & Toh, 2008). Phylogenetic trees were generated with PhyML version 3.0 using the LG substitution model plus gamma-distributed rates and four substitution rate categories (Guindon et al., 2010). The starting tree was a BIONJ tree and the type of tree improvement was subtree pruning and regrafting. Branch support was calculated using the approximate likelihood ratio test (aLRT) with a Shimodaira–Hasegawa-like (SH-like) procedure.

**Table II:** List of nuclear-encoded photosynthetic genes present in all cyanobacteria and eukaryotic phytoplankton. These genes are always nuclear-encoded, with the exception of the amoeba of the genus *Paulinella* (Fig. 2A), which has gained its plastid only very recently and independently of the event at the origin of all other known plastids (Singer et al., 2017; Yoon, Reyes-Prieto, Melkonian, & Bhattacharya, 2006).

| Gene | Pathway | Function | Copies | Non-photosynthetic homologs | References |
| --- | --- | --- | --- | --- | --- |
| *prk*<br>(phosphoribulokinase) | Calvin-Benson-Bassham cycle | phosphorylation of ribulose-5-phosphate to ribulose-1,5-bisphosphate, the RuBisCO substrate | 1 | PRKs from archaea and bacteria | (Jaffe, Castelle, Dupont, & Banfield, 2019; Kono et al., 2017) |
| *chlG*<br>(chlorophyll-a synthase) | chlorophyll-a biosynthesis | last step of chlorophyll-a biosynthesis | 1 | Prenyltransferases with UbiA domain | (Wang, Chu, Zhu, Cheng, & Yu, 2015) |
| *petH*<br>(ferredoxin-NADP<sup>+</sup> oxidoreductase) | Photosynthetic electron transport chain | last step of the linear electron flow (NADP<sup>+</sup> reduction by ferredoxin or flavodoxin) | 1-3 | -FNRs involved in nitrogen metabolism<br>-FNRs from non-photosynthetic plastids<br>-C-terminal region of benzoyl-CoA oxygenase component A (BoxA) from bacteria | (Pierella Karlusich & Carrillo, 2017; Mohamed, Zaar, Ebenau-Jehle, & Fuchs, 2001) |
| *petC* | Photosynthetic electron transport chain | Rieske subunit of the chloroplast Cyt *b<sub>6</sub>f* complex | 2-3 | Rieske proteins from mitochondria, bacteria and archaea | (Lebrun et al., 2006; Veit et al., 2016) |
| *psbO* | Photosynthetic electron transport chain | Manganese-stabilizing protein of photosystem II | 1-2 | No | (Pierella Karlusich, Ceccoli, Graña, Romero, & Carrillo, 2015) |

### Analysis of Tara Oceans datasets

*Tara* Oceans expeditions between 2009 and 2013 performed a worldwide sampling of plankton in the upper layers of the ocean (Sunagawa et al., 2020). To capture the whole size spectrum of plankton, a combination of filter membranes with different pore sizes (size-fractionation) was used to separate organisms by body size (Pesant et al. 2015). There is an inverse logarithmic relationship between plankton size and abundance (Belgrano, Allen, Enquist, & Gillooly, 2002; Pesant et al., 2015), so small size fractions represent the numerically dominant organisms in terms of cell abundance (albeit not necessarily in terms of total biovolume or biomass). Thus, the protocols consisted in the filtering of higher seawater volumes for the larger size fractions (Pesant et al., 2015). Five major organismal size fractions were collected: picoplankton (0.2-3 μm size fraction), piconanoplankton (0.8-5 μm size fraction), nanoplankton (5-20 μm size fraction), microplankton (20 to 180 μm), and mesoplankton (180 to 2000 μm). These plankton samples were leveraged to generate different molecular and optical datasets that were analysed in the current work (Table I). We exclusively used the datasets corresponding to surface samples (5 m depth).

### psbO-based community data

To use the metagenomic and metatranscriptomic read abundances of *psbO* as a proxy of phytoplankton relative cell abundance and ‘activity’, respectively, we carried out a HMMer search as stated in the previous section against the two *Tara* Oceans gene catalogues: the Ocean Microbial Reference Gene Catalog version 2 (OM-RGC.v2) covering prokaryotic and eukaryotic picoplankton (<3 µm), and the Marine Atlas of *Tara* Oceans Unigenes version 1 (MATOU.v1) covering eukaryotic plankton ranging from 0.8 to 2000 μm (Table I). The metagenomic and metatranscriptomic reads are already mapped onto both catalogs, thus we retrieved these values for those sequences obtained by our HMMer search. For the taxonomic assignment of *psbO* unigenes, we performed a phylogenetic placement of the translated sequences on the PsbO protein reference phylogenetic tree described in the previous section. For parallelization of the task, a set of 50 unigenes were translated and the PsbO specific Pfam PF01716 region was retrieved for the analysis in the following way. First, they were aligned against the reference alignment described in the previous section using the option --add of MAFFT version 6 with the G-INS-I strategy (Katoh and Toh 2008). The resulting alignment was used for building a phylogeny with PhyML version 3.0 as described above (Guindon et al., 2010). The sequences were classified using the APE library in R (Paradis & Schliep, 2019) according to their grouping in monophyletic branches of statistical support >0.7 with reference sequences of the same taxonomic group.

Due to challenges of assembling eukaryotic genomes from complex metagenomes, the MATOU-v1 catalog only contains sequences assembled from poly-A-tailed RNA (Alberti et al., 2017; Carradec et al., 2018), which biases against prokaryotic sequences. To determine the structure of the whole phytoplankton community (including both cyanobacteria and eukaryotic phytoplankton), we aligned all the metagenomic reads from *Tara* Oceans to a curated database of *psbO* sequences (described below; see also Table I). The analysis was carried out using the bwa tool version 0.7.4 (H. Li & Durbin, 2009) with the following parameters: – minReadSize 70-identity 80-alignment 80-complexityPercent 75-complexityNumber 30. Abundance values were expressed in rpkm (reads per kilobase covered per million of mapped reads).

In general, the rpkm values for the different taxa under study were converted to percentage of (either total or eukaryotic) phytoplankton. However, for a specific analysis the *psbO* rpkm values were normalized by those values form single-copy housekeeping genes: by bacterial *recA* (Sunagawa et al. 2013) to estimate the contribution of cyanobacteria in the bacterioplankton, or by the average abundance of 25 genes encoding ribosomal proteins (Ciccarelli et al. 2006; Carradec et al. 2018) to estimate the contribution of phytoplankton among eukaryotes. The abundance values for *recA* were retrieved from a previous work (Pierella Karlusich et al. 2021) while the ribosomal proteins were recovered from the MATOU-v1 and OMRGC-v2 abundance tables.

### rRNA gene-based community data

We used two different datasets generated by *Tara* Oceans for “traditional” DNA-based methods: 16S rRNA gene miTags (size fraction 0.2-3 µm) and 18S rRNA gene (V9 region) metabarcoding (sizes fractions 0.8-5, 5,20, 20-180, 180-2000 µm) (Table I). We extracted the relative abundances for the 726 Operational Taxonomic Units (OTUs) assigned to picophytoplankton (cyanobacteria and chloroplasts) from the 16S miTags and the 31,930 OTUs assigned to eukaryotic phytoplankton from the V9-18S metabarcoding data. The read abundances were expressed as relative abundance (%) in relation to the picophytoplankton community for 16S miTags, and in relation to eukaryotic phytoplankton for V9-18S metabarcoding.

The assignations of the 16S and V9-18S rRNA sequences to phytoplankton were based on literature and expert information and included photosynthetic dinoflagellates and chrysophytes when their taxonomic resolution was sufficient to match known photosynthetic lineages. A full description of the 18S taxonomic classification procedure is at http://taraoceans.sb-roscoff.fr/EukDiv/ and the last version of the trait reference database used in the current work is available at https://zenodo.org/record/3768510#.YM4ny3UzbuE. In the case of 16S miTags, the taxonomic assignment was improved by building a phylogenetic tree with the 16S miTags sequences and a curated set of references from NCBI and MMETSP. Sequences were aligned using MAFFT v7.0 (Katoh & Standley, 2013) with --auto setting option and then trimmed using trimal with the -gt 0.5 and -gt 0.8 settings, and the resulted alignment was used for tree building using RAxML v8 (Stamatakis, 2014) (100 bootstrap replicates, GTRCAT substitution model).

### Optical-based community data

We also used quantitative optical data generated by *Tara* Oceans (Table I), where cell abundance is assumed to be more accurate and less biased, and additional features such as biovolume can be determined. The datasets cover: flow cytometry for picoplankton, confocal microscopy for 5–20 μm size fraction, and light microscopy for 20-180 µm size fraction.

Flow cytometry counts were determined on 1 ml replicated seawater samples filtered through 200 μm that were fixed with cold 25% glutaraldehyde (final concentration 0.125%) and stored at -80°C until analysis. Details about the procedure can be found in (Gasol & Morán, 2015; Hingamp et al., 2013; Pierella Karlusich et al., 2021). The cell biovolume was calculated using the equation of (Calvo-Díaz & Morán, 2006) on the bead-standardized side scatter of the populations and considering cells to be spherical.

Quantitative confocal microscopy was performed using environmental High Content Fluorescence Microscopy (eHCFM) (Colin et al., 2017). Briefly, samples were fixed with 10% monomeric formaldehyde (1 % final concentration) buffered at pH 7.5 and 500 μl EM grade glutaraldehyde (0.25% final concentration) and kept at 4 °C until analysis. Sample collection and preparation as well imaging acquisition is described in (Colin et al., 2017). The 5–20 μm size fraction has been classified at a coarse taxonomic level (with an estimated accuracy of 93.8% at the phylum or class level), into diatoms, dinoflagellates, haptophytes, and other/unclassified eukaryotic phytoplankton (Colin et al., 2017). We used the major and minor axis of every image to calculate their ellipsoidal equivalent biovolume. The 20-180 µm size fraction is also available, but the curated taxonomic annotation is limited to symbiotic (*Richelia*, *Calothrix*) and colony-forming (*Trichodesmium*) nitrogen-fixing cyanobacteria (Pierella Karlusich et al. 2021), which were also used in the current work.

For light microscopy, three ml of each sample (from 20-180 µm size fractions) were placed in an Utermöhl chamber with a drop of calcofluor dye (1:100,000) which stains cellulose, thus allowing to better detect and identify dinoflagellates. Cells falling in 2 or 4 transects of the chamber were identified and enumerated using an inverted light microscope (Carl Zeiss Axiophot200) at 400x magnification.

To be compared with the molecular data, the optical data were expressed as relative abundance (%). In the case of flow cytometry, as % over the total number of cells counted as picophytoplankton (*Prochlocococcus* + *Synechococcus* + eukaryotic picophytoplankton). In the case of confocal and optical microscopy, as % over the total number of cells counted as eukaryotic phytoplankton.

### psbO database generation

We compiled, curated and annotated a database of >18,000 unique *psbO* sequences covering cyanobacteria, photosynthetic protists, macroalgae and land plants (Figure S1). It includes sequences retrieved from IMG, NCBI, MMETSP and other sequenced genomes and transcriptomes from cultured isolates, as well as from the environmental sequence catalogs from Global Ocean Sampling (Rusch et al., 2007) and *Tara* Oceans (Carradec et al., 2018; Delmont et al., 2020, 2021; Salazar et al., 2019). The database can be downloaded from the EMBL-EBI repository BioStudies (www.ebi.ac.uk/biostudies) under accession S-BSST659. We expect to maintain it updated to facilitate its incorporation in molecular-based surveys.

### Plotting and statistical analysis

Graphical analyses were carried out in R language (http://www.r-project.org/) using *ggplot2* (Wickham, 2016) and treemaps were generated with *treemap*. Maps were generated with borders function in *ggplot2* and *geom_point* function for bubbles or *scatterpie* package for pie charts (Yu, 2018). Spearman’s rho correlation coefficients and p-values were calculated using the *cor.test* function of the *stats* package. Shannon diversity indexes were calculated using the *vegan* package (Oksanen et al. 2020). Intra- and inter-specific genetic distances were calculated in MEGAX (Kumar, Stecher, Li, Knyaz, & Tamura, 2018) using the Maximum Composite Likelihood model.

## Results

### Search for phytoplankton marker genes

We first analysed transcriptomes and nuclear and plastid genomes derived from cultured strains to inventory photosynthetic genes in relation to their genome location (nuclear- vs plastid-encoded) and taxonomic prevalence (core vs non-core, i.e., present in all phototrophs or not) (see Methods; Figure 1A). Among the plastid-encoded genes, we identified phytoplankton marker genes previously used in environmental surveys, such as *psbA* (Man-Aharonovich et al., 2010b; Zeidner, Preston, Delong, Massana, Post, Scanlan, & Béjà, 2003), *rbcL* (nuclear encoded in dinoflagellates containing peridinin) (Paul et al., 2000) and *petB* (Farrant et al., 2016) (Figure 1A).

**Figure 1:**
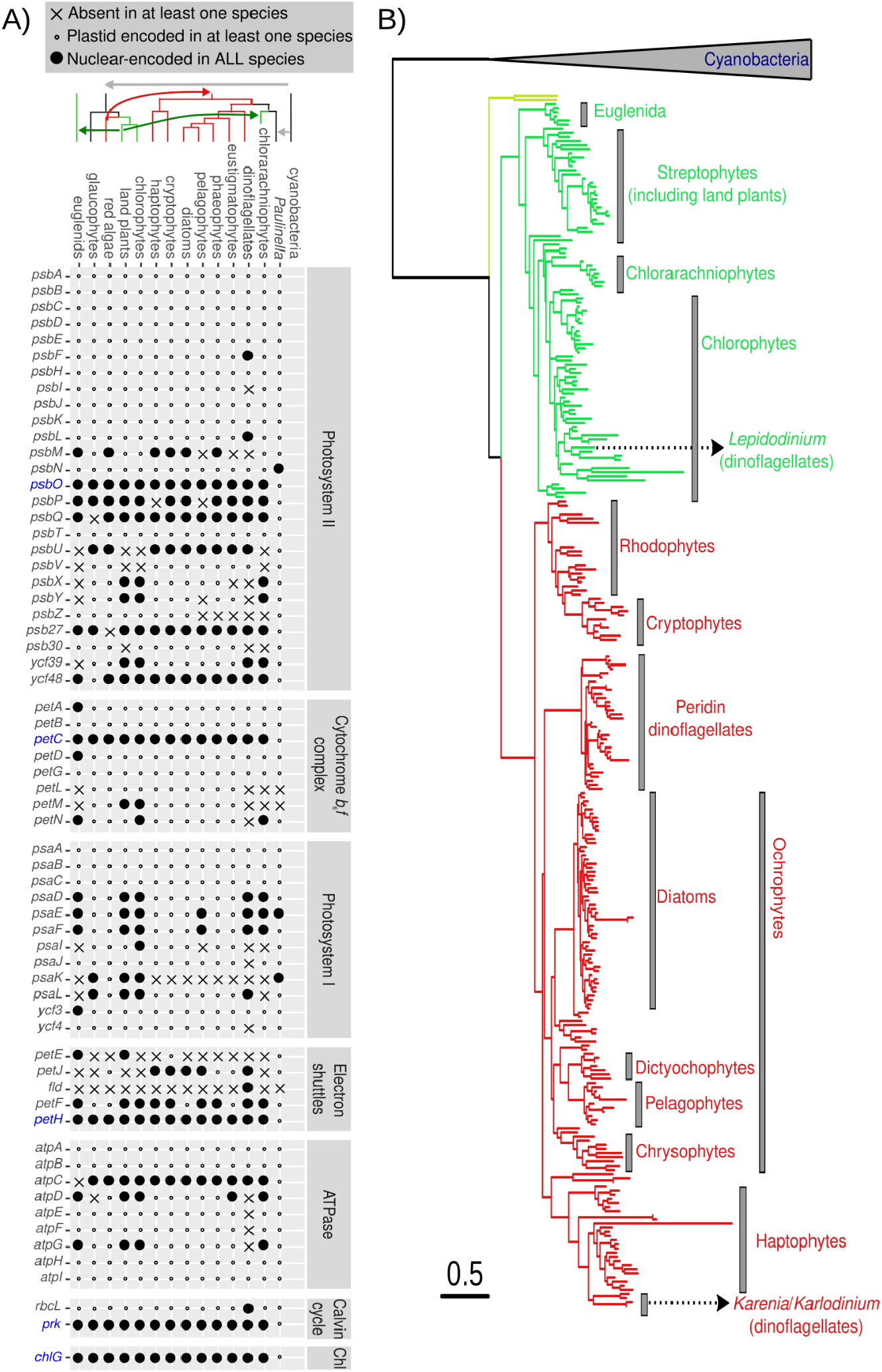
Identification of nuclear-encoded core photosynthetic gene marker candidates. A) Presence and location of the genes encoding proteins involved in photosynthesis. The genes found to be core and nuclear-encoded are labelled in blue. The only exception is the amoeba of the genus *Paulinella*, which has gained its plastid very recently and independently of the event at the origin of all other known plastids, thus still retaining these genes in its plastid genome (Yoon et al. 2006; Singer et al. 2017). B) Phylogeny of PsbO protein. Translated sequences from genomes and transcriptomes of cultured phytoplanktonic species were used for the phylogeny reconstruction. The scale bar indicates the number of expected amino acid substitutions per site per unit of branch length.

Among the nuclear-encoded genes, we retrieved some which are non-core, such as those encoding flavodoxin (*fld*) and plastocyanin (*petE*), but also five core genes (Figure 1A; Table II). These five genes are present in low-copy number and encode components of the photosynthetic electron transport chain (*psbO*, *petC* and *petH*), the carbon fixation pathway (*prk*) or chlorophyll biosynthesis (*chlG*) (Table II). The absence of non-photosynthetic homologs is a unique characteristic of *psbO* (Table II and Figures S2-S6), reflecting its essential role in photosynthetic oxygen evolution, and a clear advantage for its use as a marker gene for phytoplankton. Previous studies of secondarily non-photosynthetic eukaryotes have marked its presence or absence as being an effectively universal predictor of photosynthetic potential (Dorrell et al., 2019). Its phylogeny additionally reflects the evolutionary history of endosymbiosis (Figure 1B; Pierella Karlusich et al. 2015), with few or no post-endosymbiotic horizontal replacements known, so we focused on this gene for the analysis of environmental samples.

Although no global barcoding gap (i.e., a distance threshold set for all species) was detected when checking intra- vs inter-specific divergences for eukaryotic phytoplankton based on *psbO*, it was neither observed with the V9 region of the traditional marker 18S rRNA gene (Figure S7). This absence does not necessarily preclude specimen identification, which relies upon the presence of a ‘local’ barcoding gap (i.e., a query sequence being closer to a conspecific sequence than a different species), rather than the ‘global’ barcoding gap (i.e., a distance threshold set for all species) that is required for species discovery (Collins & Cruickshank, 2013).

We retrieved the *psbO* sequences from the two *Tara* Oceans gene catalogs (the picoplankton catalog OM-RGC.v2 and the eukaryotic catalog MATOU.v1; see Methods and Table I). A total of 307 distinct sequences were identified in OM-RGC.v2 (202 from *Prochlorococcus*, 79 from *Synechoccocus* and 26 from eukaryotic picophytoplankton), with an average length for the conserved coding region of 473 base pairs and a range between 94 and 733 base pairs. A total of 10,646 sequences from eukaryotic phytoplankton were retrieved from MATOU.v1, with an average length for the conserved coding region of 385 base pairs and a range between 66 and 784 base pairs. The analyses of the metagenomic and metatranscriptomic read abundances of these sequences are presented in the following sections.

### Marine phytoplankton community structure based on psbO shows remarkable differences with the traditional molecular approaches

The abundance and diversity of phytoplankton was first examined in *Tara* Oceans samples by focusing on the traditional marker genes coding for the small subunit of rRNA (16S for prokaryotes and plastids, 18S for eukaryotes) in the different size-fractionated samples. We focused exclusively on the phytoplankton signal of these datasets, despite the uncertainties in assigning photosynthesis capacity in groups such as dinoflagellates and chrysophytes (this is evaluated in one of the next sections).

Based on 16S miTag read abundance among picophytoplankton (0.2-3 µm), the picocyanobacteria *Prochlorococcus* and *Synechococcus* were prevalent, while ∼60% of the average read abundance was attributed to eukaryotic photosynthetic taxa such as haptophytes, chlorophytes, pelagophytes, dictyochophytes, chrysophytes, cryptophytes and diatoms (Figure 2A). In the larger size fractions, based on the V9-18S region metabarcoding reads, diatoms and dinoflagellates were the most frequent among eukaryotic phototrophs, especially in the 5–20 μm and 20–180 μm size fractions (Figure 2B). In the 180-2000 μm fraction, diatoms and dinoflagellates were still abundant, due to the presence of large diameter cells (*Tripos*, *Pyrocystis*), chain-forming (e.g., *Chaetoceros*, *Fragilaria*) or epizoic (e.g., *Pseudohimantidium*) species, without discarding that smaller species may be retained in samples of this size fraction due to net clogging or within herbivorous guts and faecal pellets. Relative abundance in the smaller 0.8–5 μm size fraction was much more homogeneously distributed between the different groups.

**Figure 2:**
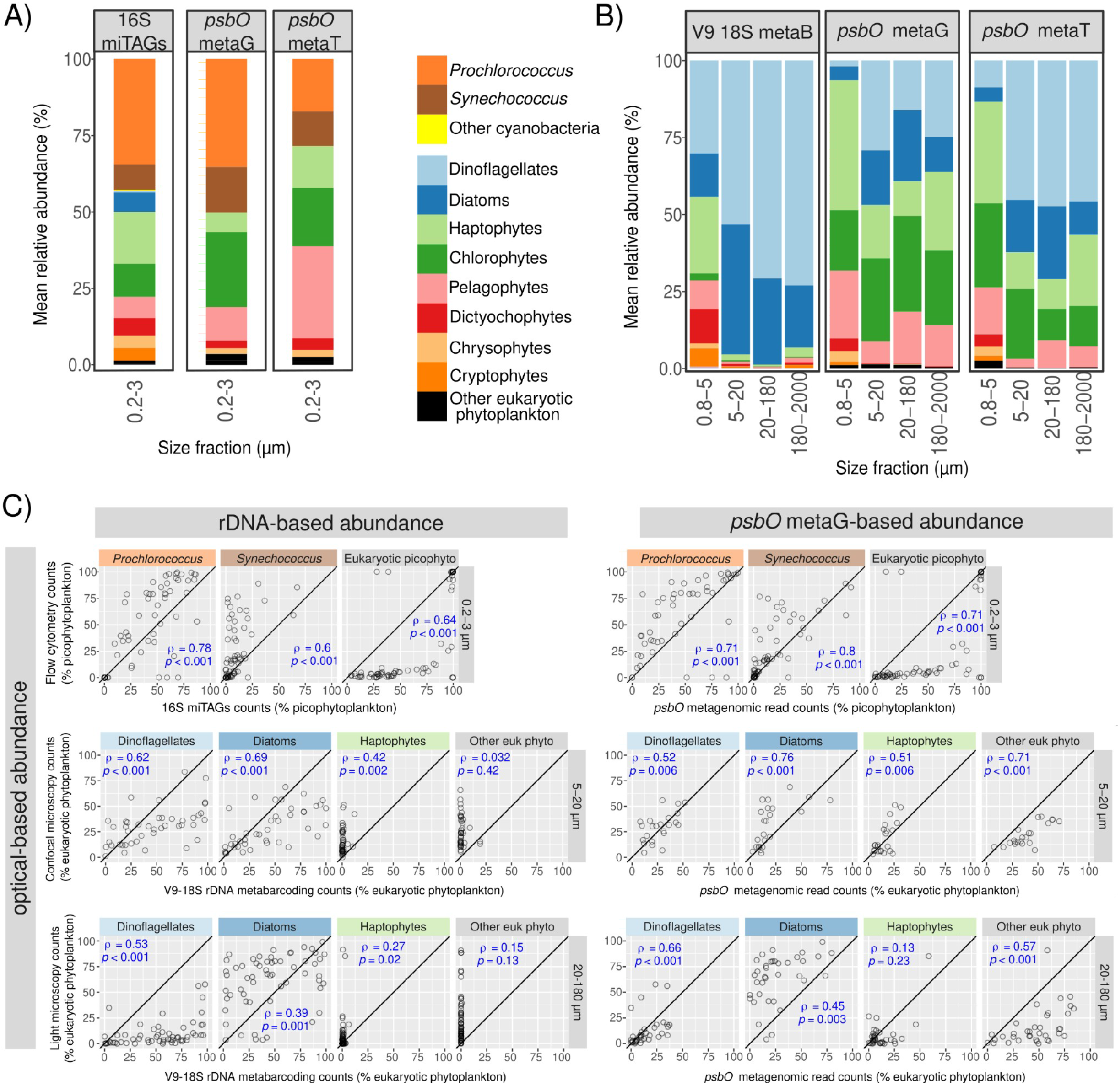
Congruence in relative abundances of the main phytoplankton groups based on different gene markers. A-B) Average relative abundances for all surface samples of each size fraction using different marker genes. In A, picocyanobacteria and eukaryotic picophytoplankton (0.2-3 μm) were analysed using 16S rRNA gene miTags and the metagenomic and metatranscriptomic read abundances for *psbO*. In B, eukaryotic phytoplankton was analysed in larger size fractions using V9-18S rRNA gene amplicons and the metagenomic and (polyA-derived) metatranscriptomic *psbO* read abundances. C) Correlations between relative abundances of different phytoplankton groups obtained with optical versus DNA-based methodologies. In the upper panel, 16S rRNA gene miTags and *psbO*-based relative abundances in picophytoplankton were compared with flow cytometry counts (values displayed as % total abundance of picophytoplankton). In the middle and lower panels, V9-18S rRNA gene metabarcoding and metagenomic *psbO* relative abundances were compared with confocal microscopy counts from size fraction 5-20 μm and light microscopy counts from size fraction 20-180 µm (values displayed as % total abundance of eukaryotic phytoplankton). It is worth mentioning that the molecular and microscopy data were generated from the same samples, while there were differences between molecular data of 0.2-3 µm size fraction and flow cytometry data (see Methods). Axes are in the same scale and the diagonal line corresponds to a 1:1 slope. Spearman’s rho correlation coefficients and p-values are displayed. The correlations of relative abundances between metatranscriptomic *psbO* reads and optical methods are shown in Figure S11.

For *psbO*-based methods, we found that metagenomic and metatranscriptomic reads from *Synechococcus*, *Prochlorococcus*, pelagophytes, chlorophytes and haptophytes to be dominant among picophytoplankton (0.2-3 µm), along with dictyochophytes and chrysophytes (Figure 2A). In the larger size fractions, haptophytes, chlorophytes and pelagophytes clearly dominated the eukaryotic phytoplankton in the 0.8-5 μm size fraction, whereas diatoms and dinoflagellates were more abundant in the three larger size ranges (5-20 μm, 20-180 μm, 180-2000 μm), although haptophytes, chlorophytes and pelagophytes were also detected in large quantities (Figure 2B). The potential cyanobacteria present in these large size fractions are presented later in another section due to the need to bypass the sequences assembled from poly-A-tailed RNA for analysing prokaryotes (see Methods and Table I).

We noted some differences in *psbO* read counts between metagenomic and metatranscriptomic datasets. In the case of picophytoplankton, *Prochlorococcus* was enriched in metagenomes in comparison to (total RNA) metatranscriptomes (likely due to low transcript content needs for their low cell volume), the opposite occurred for pelagophytes and haptophytes, whereas no major changes were observed for *Synechococcus* and chlorophytes (Figures 2A and S8A). In the case of larger photosynthetic protists, dinoflagellates were highly abundant at the (polyA) transcript level in comparison to gene abundance (probably they blanket overtranscribe genes as they predominantly perform post-transcriptional gene regulation (Roy, Jagus, and Morse 2018; Cohen et al. 2021)), the opposite was observed for pelagophytes and chlorophytes (in this latter taxon only in the 20-180 and 180-2000 μm size ranges), whereas no major shifts were apparent for diatoms and haptophytes (Figures 2B and S8B).

The taxonomic abundance patterns based on *psbO* showed some differences with those from 16S miTags of 0.2-3 µm size fraction, but exhibited remarkable differences with those based on V9-18S metabarcoding of the large size fractions (Figures 2A-B and S9). When compared with the 16S miTags, no major changes were detected for *Prochlorococcus*, whereas the average *psbO* metagenomic contribution increased for *Synechococcus* (from ∼8% to ∼14%), at the expense of decreasing eukaryotic picoplankton contribution (from 57% to ∼50%), which is expected due to the fact that the 16S rRNA is a plastid-encoded gene in eukaryotes. When we compared *psbO* with V9-18S metabarcoding, the differences were very significant. In the 0.8-5 μm size fraction, diatoms and dinoflagellates accounted for just ∼6% of average *psbO* metagenomic read abundance but for ∼44% of 18S reads assigned to phytoplankton. In the three larger size ranges (5-20 μm, 20-180 μm and 180-2000 μm), they accounted for 37-47% of average *psbO* metagenomic read abundance, but for >90% of average 18S read abundance. The 18S read abundance was extremely low for haptophytes, chlorophytes and pelagophytes in these three size fractions (<7% average 18S read abundance). When we compared the metatranscriptomic profile, it was more similar to the profile obtained with metagenomes than to that obtained with V9-18S metabarcoding (Figure 2B).

### Comparison with imaging dataset suggests that psbO is a robust marker gene for estimating relative cell abundance of phytoplankton from metagenomes

To assess the accuracy of the *psbO* gene for determining phytoplankton cell relative abundances, we carried out comparative analyses with imaging datasets. For the 0.2-3 µm size fraction, we compared relative abundances based on 16S and *psbO* counts with those inferred from flow cytometry (Figure 2C). Both genes were found to correlate well with flow cytometry. Although the correlations for eukaryotic picophytoplankton were strong (Spearman’s rho=0.64-0.71, p-value<<<0.001), the relationships were not linear and picoeukaryotes appeared at much higher relative abundances in metagenomes than in flow cytometry. This is consistent with the fact that flow cytometry can count cells of up to 10-20 µm diameter and was performed on seawater aliquots pre-filtered through a 200-μm mesh (see Methods), whereas DNA isolation of picoplankton was carried out on seawater volumes mainly filtered through 3-μm pore sizes. When we discarded eukaryotes to focus only on the ratio *Synechococcus* / (*Synechococcus* + *Prochlorococcus*) (Figure S10), flow cytometry data shows a linear relashiopship with *psbO* metagenomic reads, while 16S miTags reads underestimated *Synechococcus* and the opposite occurred for *psbO* metatranscriptomic reads. In addition, the highest correlation with flow cytometry data occurred with the *psbO* metagenomic counts (Spearman’s rho=0.92, 0.90 and 0.75, p<<<0.001, for *psbO* metagenomic reads, *psbO* metatranscriptomic reads and 16S miTags, respectively).

For the 5–20 μm size fraction, the relative abundance of eukaryotic photosynthetic organisms was determined by cell counts using high-throughput confocal microscopy. We compared these results with the proportions based on V9-18S metabarcoding and *psbO* metagenomic reads (Figure 2D). The metabarcoding data for dinoflagellates and diatoms were in good agreement with the microscopy but it clearly underestimated the relative abundance of haptophytes and other eukaryotic phytoplankton. Regarding *psbO*, the metagenomic relative abundances were in stronger agreement with the microscopy counts for the four defined phytoplankton groups (Figure 2D). Therefore, in the 5–20 μm size fraction, diatoms and dinoflagellates displayed robust patterns of relative abundance using either V9-18S metabarcoding or *psbO* metagenomic counts, while haptophytes and the other groups were better described by *psbO*.

In the 20-180 μm size fraction, the relative abundance of eukaryotic phytoplankton was determined by light microscopy. Again, the metabarcoding data for dinoflagellates and diatoms were in good agreement with the microscopy data but clearly underestimated the relative abundance of haptophytes and other eukaryotic phytoplankton groups (Figure 2D). The metagenomic read relative abundances of *psbO* were in stronger agreement with the microscopy counts for the four defined phytoplankton groups, although the correlation with haptophytes was weaker (Figure 2D). Therefore, in the 20-180 μm size fraction, diatoms and dinoflagellates displayed robust patterns of relative abundance using either V9-18S metabarcoding or *psbO* metagenomic counts, while haptophytes were weakly described by both methods and the other groups were much better described by *psbO*.

We also compared the relative abundances based on optical methods against those based on *psbO* metatranscriptomic reads, and in general we observed good agreement (Figure S11). Some phytoplankton groups displayed stronger correlations against optical methods using metatranscriptomic *psbO* counts than 16S miTAGs (e.g., *Synechococcus*) or V9-18S metabarcoding (e.g., other eukaryotic phytoplankton in the 5-20 µm) (Figures 2D and S11). However, the consistency in relative abundance of *psbO* reads with optical methods was always better for metagenomes than for metatranscriptomes (Figures 2D and S11).

### Comparison with optical-based biovolume suggests that neither psbO nor rRNA genes are good proxies for estimating relative proportion of biovolume

We also compared the relative read abundances of the different marker genes against the proportional biovolumes for each taxon (Figure 3). Although the copy number of rRNA marker genes was previously proposed as a proxy of cell biovolume, the correlation of biovolume against rRNA gene relative abundances was not stronger than those against *psbO* (Figures 3 and S12). The relative read abundances for *Prochlorococcus* and eukaryotic picophytoplankton based either on 16S rRNA gene or *psbO* were higher than their proportional biovolumes in the same samples, while the opposite was the case for *Synechococcus.* In the 5-20 µm size fraction, the biovolume proportion for haptophytes was clearly described by their *psbO* relative abundance, while their V9-18S rRNA gene reads were very low in relation to their biovolume. Both V9-18S rRNA gene and *psbO* reads were correlated with the relative biovolume for diatoms and dinoflagellates, but for V9-18S rRNA gene the data points were somewhat scattered and for *psbO* the relative abundances for the reads were higher in relation to their biovolume. As the biovolume of other taxa was very low, their proportion of *psbO* reads was much higher than the corresponding biovolume fraction, whereas there was no correlation between V9-18S and biovolume.

**Figure 3:**
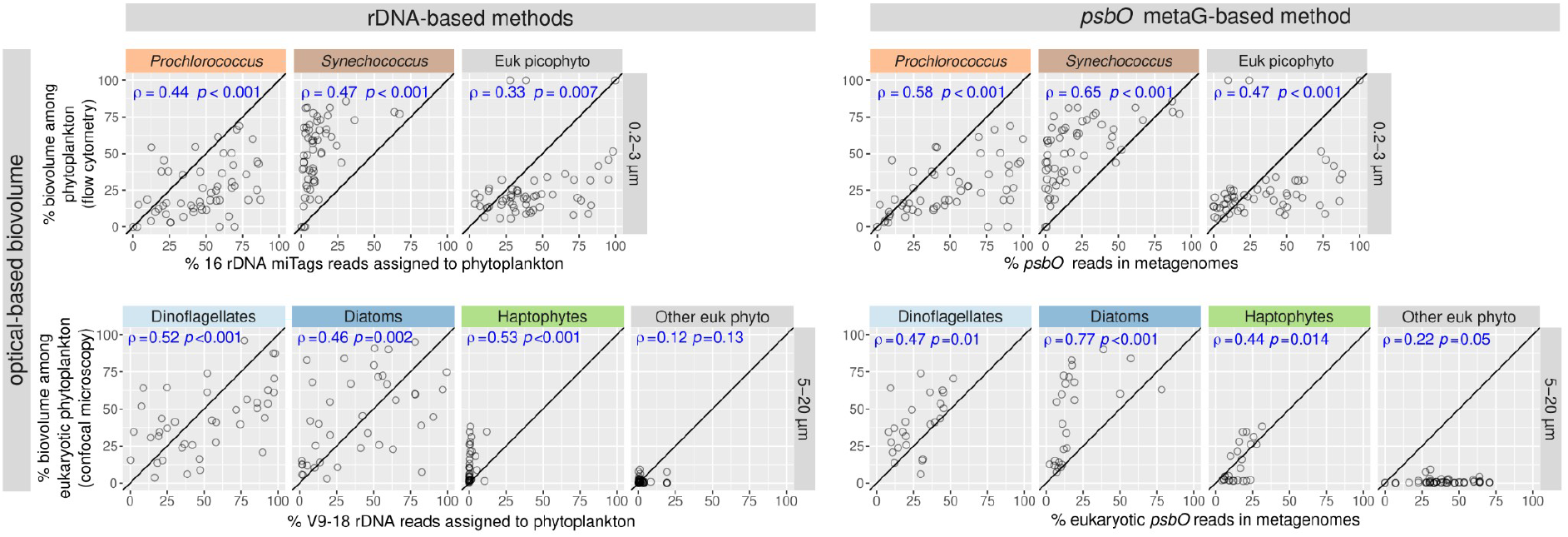
Correlation between relative biovolume (based on optical methods) and relative abundances based on different molecular methodologies. The upper panels show the correlations for picophytoplankton (size fraction 0.2-3 µm). The vertical axis corresponds to the relative biovolume based on flow cytometry (values displayed as % total biovolume of picophytoplankton), while the horizontal axis corresponds to relative read abundance based on molecular methods: 16S miTAGs (left upper panel) and *psbO* metagenomic counts (right upper panel). The lower panels show the correlations for nanophytoplankton (size fraction 5-20 μm). The vertical axis corresponds to the relative biovolume based on confocal microscopy quantification (values displayed as % total abundance of eukaryotic phytoplankton), while the horizontal axis corresponds to relative read abundance based on molecular methods: V9-18S rRNA gene metabarcoding (left lower panel) and eukaryotic *psbO* metagenomic counts (right bottom panel). Spearman correlation coefficients and p-values are displayed in blue. Axis are in the same scale and the diagonal line corresponds to a 1:1 slope.

### Diversity analysis: Shannon-index is robust to the biases introduced by the traditional molecular methods

We further analysed whether our method improved the widely used Shannon index, a diversity index that relates monotonically to species richness but differs in that it downweights rare species, whose numbers are highly sensitive to undersampling and molecular artefacts (Calderón-Sanou, Münkemüller, Boyer, Zinger, & Thuiller, 2020). We found a strong correlation between Shannon values for eukaryotic phytoplankton defined either by 18S rRNA gene metabarcoding or by *psbO* metagenomics or metatranscriptomics (Figure 4). This is in agreement with previous reports showing no major effects of 16S rRNA gene copy number variation on the Shannon index of bacterial communities (Ibarbalz et al., 2019; Milanese et al., 2019). These results illustrate that not all subsequent analyses are affected by the biases introduced by traditional molecular methods.

**Figure 4:**
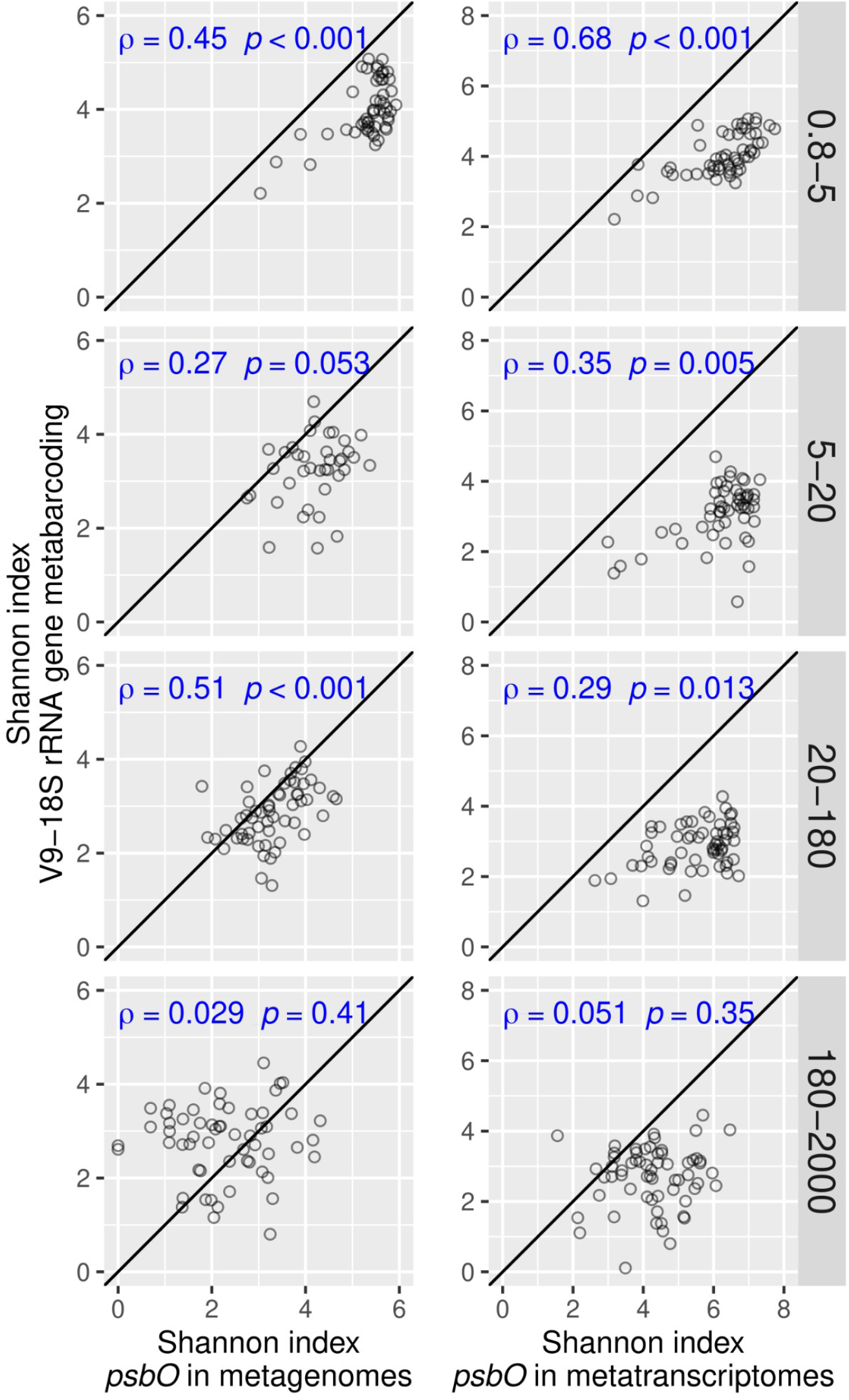
Correlation between the Shannon values derived from different molecular methods for eukaryotic phytoplankton communities. The values derived from *psbO* metagenomics (left) and metatranscriptomics (right) were compared with those derived from V9-18S rRNA gene metabarcoding. Spearman correlation coefficients and p-values are displayed in blue. Axis are in the same scale and the diagonal line corresponds to a 1:1 slope.

### Combining housekeeping and photosynthetic marker genes improves estimates of the distribution and abundance of phototrophs in a given taxonomic group

To evaluate the uncertainties when inferring the photosynthesis trait using the taxonomy obtained from a non-photosynthetic marker gene, we analysed the 18S V9 OTUs assigned to dinoflagellates and found that most of their reads cannot be reliably classified as corresponding or not to a photosynthetic taxon (Figure 5A), especially for those OTUs whose taxonomic affiliation is “unknown dinoflagellate” (Figure S13). The uncertainty was especially significant in the 0.8-5 µm size fraction, where on average the ∼80% of the total dinoflagellate read abundance remained unclassified (Figures 5A and S13).

**Figure 5:**
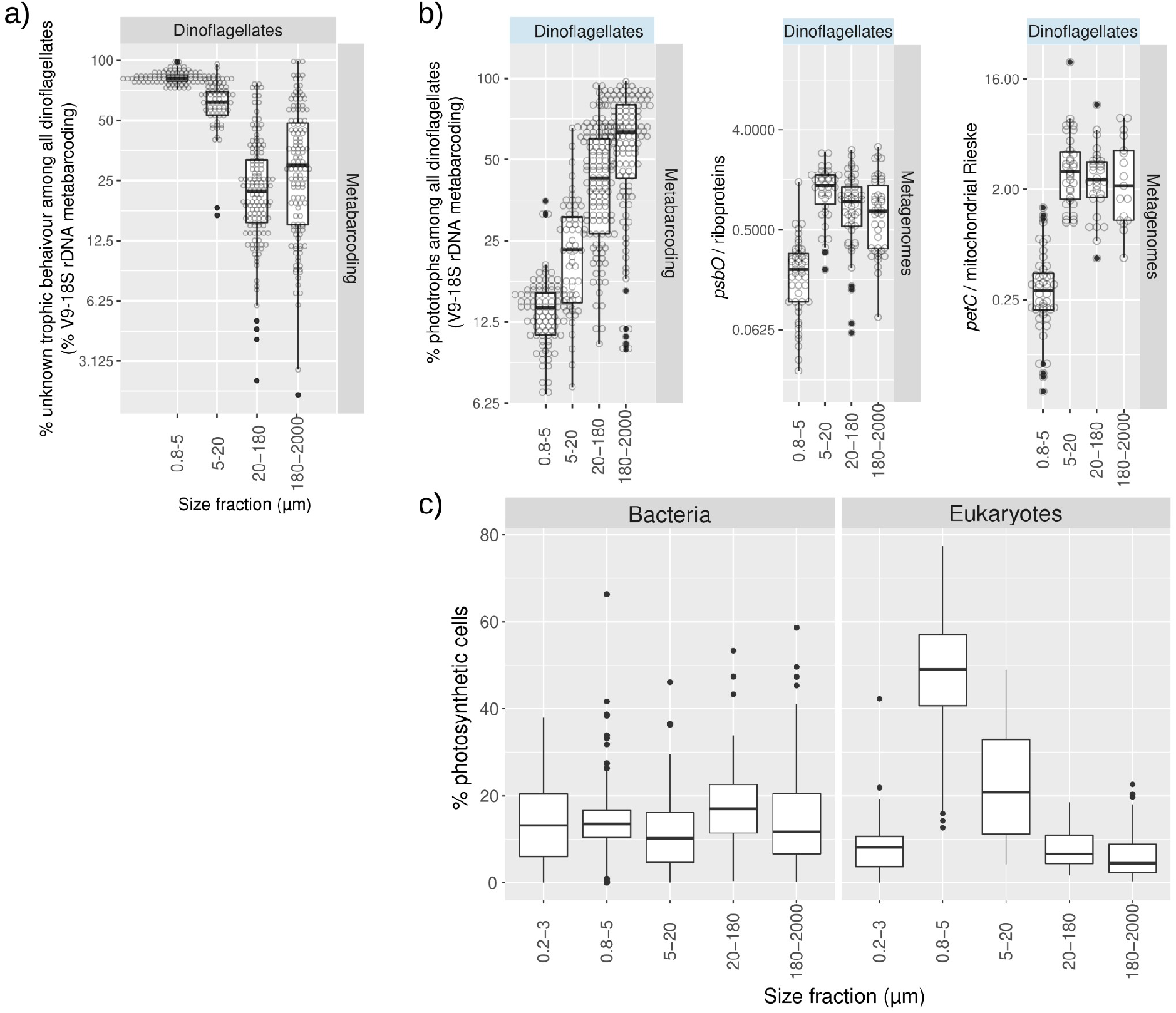
Variations in the abundance of phototrophs *vs* heterotrophs across size fractions. A) Read abundance of V9-18S rRNA gene metabarcoding assigned to dinoflagellates of unknown capacity for photosynthesis. B) Relative abundance of phototrophs among dinoflagellates based on different molecular methods. The first panel corresponds to the trait classification of V9-18S rRNA gene metabarcodes based on the literature (a description of the trait classification can be found at http://taraoceans.sb-roscoff.fr/EukDiv/ and the trait reference database is available at https://zenodo.org/record/3768951#.YM4odnUzbuE). The second and third panels correspond to the ratio of metagenomic counts of photosynthetic *vs* housekeeping single-copy nuclear-encoded genes: *psbO* vs genes coding for ribosomal proteins, and the genes coding for the Rieske subunits of the Cyt bc-type complexes from chloroplasts and mitochondria (i.e., *petC* and its mitochondrial homolog). C) Relative abundance of phototrophs among bacterial and eukaryotic plankton across size fractions. The values were determined by the ratio of metagenomic counts of the single-copy marker genes of photosynthesis (i.e., *psbO*) and housekeeping metabolism (i.e., *recA* for bacteria and genes encoding ribosomal proteins for eukaryotes).

Therefore, besides finding a more relevant marker gene for phytoplankton, we also propose combining it with established single-copy housekeeping genes (i.e., *recA* for bacteria (Sunagawa et al., 2013) and genes encoding ribosomal proteins for eukaryotes (Carradec et al., 2018; Ciccarelli et al., 2006)), to estimate the fraction of photosynthetic members in a given community or within a specific clade. In the case of eukaryotes, a set of genes of interest for this aim are *petC* and its mitochondrial homologs (i.e., the nuclear-encoded genes for the Rieske subunits of the Cyt *bc*-type complexes from chloroplasts and mitochondria) (Table II and Figure S5). As an example, we analysed the distribution of phototrophy across size fractions among the eukaryotic groups under study. As expected, it did not reveal any differences for diatoms, haptophytes, chlorophytes or pelagophytes (Figure S14), reflecting the relative paucity of described secondarily non-photosynthetic members of these groups. Instead, for dinoflagellates we observed a significant proportion of non-photosynthetic lineages in the 0.8-5 size-fraction in comparison with the other sizes ranges, which were also shown by the V9-18S rRNA gene metabarcoding method (Figures 5B, S13 and S14). However, whereas the metabarcoding data showed a dramatic increase in phototrophs towards the larger size classes of dinoflagellates, the metagenomic analysis showed similar levels between the three larger size fractions (5-20 µm, 20-180 µm, 180-2000 µm) (Figure 5B). These different patterns between the two marker genes might be explained by differences in the unknown trait assignment of the 18S rRNA gene barcodes and/or in the 18S rRNA gene copy number (e.g., higher copy number in photosynthetic species in larger size fractions).

The approach suggested can be applied to unveil variation of phototrophs in whole plankton communities, including both bacteria and eukaryotes. In order to do so, we mapped the metagenomic reads against our comprehensive catalog of *psbO* sequences (Figure S1). The highest proportion of phytoplankton among eukaryotes was observed in the 0.8-5 µm size fraction, followed by the 5-20 µm size-fraction, while the lowest value was found in the 180-2000 µm size range (Figure 5C), where copepods are prevalent (considered one of the most abundant animals on the planet). Surprisingly, the percentage of phototrophs among bacterioplankton did not vary across size fractions (10-15 % on average; see next section). In the 0.2-3 µm size fraction, very similar values were detected by 16S miTags, but when comparing both molecular methods with flow cytometry, the *psbO*/*recA* ratio was better correlated to flow cytometry (Spearman’s rho of 0.82 vs 0.91, p<0.001, and a closer 1:1 relationship) (Figure S15).

### Trans-domain comparison reveals unexpected abundance of picocyanobacteria in large size fractions

To further examine the distribution of both prokaryotic and eukaryotic phytoplankton across the whole size spectrum, we continued the analysis of the mapped metagenomic reads against our catalogue of *psbO* sequences (Figure S1). We observed a high abundance of cyanobacteria in the large size fractions in relation to the eukaryotic phytoplankton (Figure 6A). The nitrogen-fixers *Trichodesmium* and *Richelia*/*Calothrix* were found principally in the 20-180 and 180-2000 μm size fractions (Figure 6A), which is expected as the former forms filaments and colonies while the second group are symbionts of certain diatoms (Figures 7B-D). These genera were recently quantified in the high-throughput confocal microscopy dataset from the 20-180 µm size fraction (Pierella Karlusich et al., 2021). We therefore checked the correlations of these data with the *psbO* determinations and found them to be very strongly related (Figure 6G).

**Figure 6:**
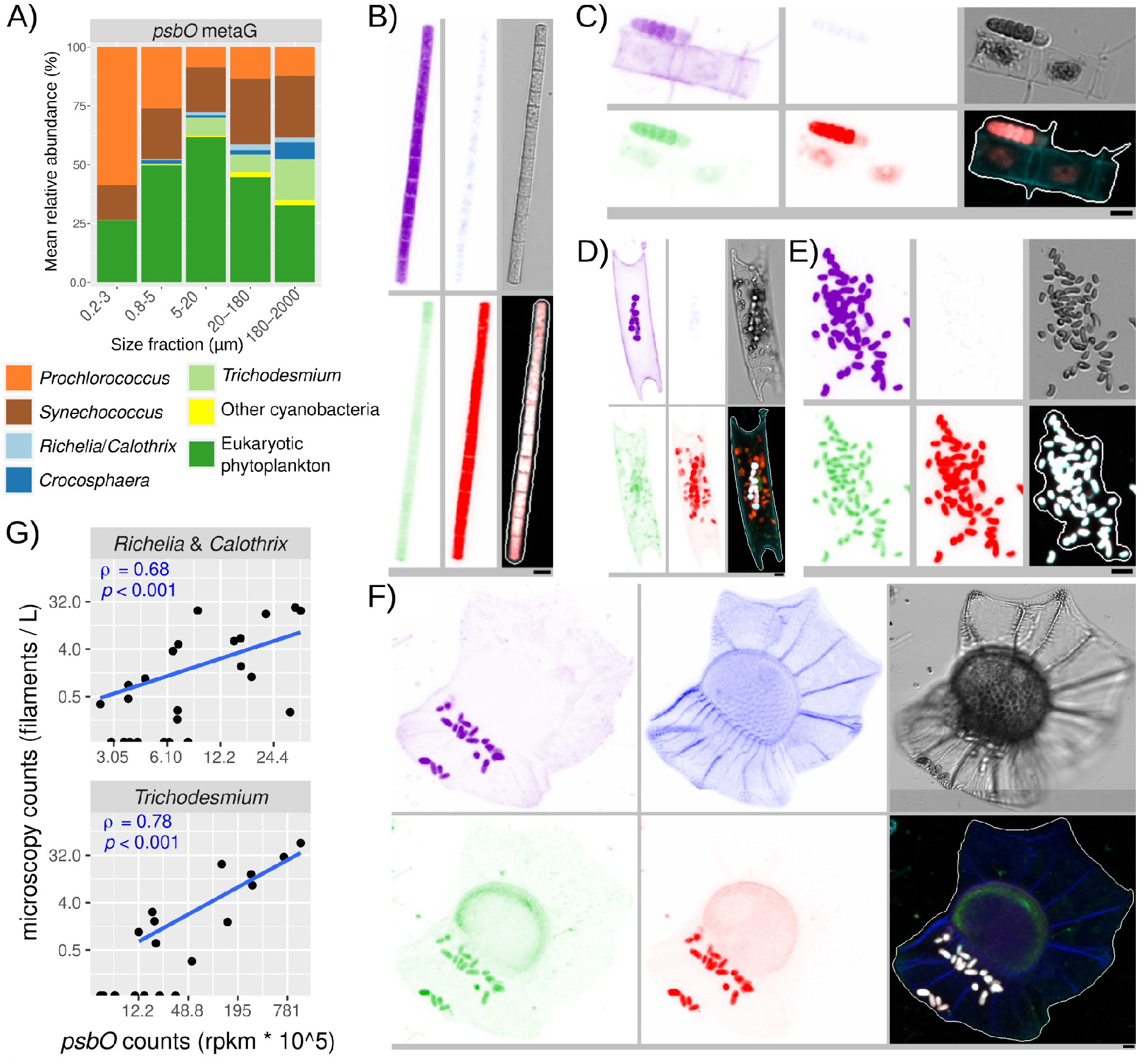
Prokaryotic and eukaryotic phytoplankton community structure across the entire plankton size spectrum. A) Average relative cell abundance of phototrophs across all metagenomes based on *psbO* metagenomic reads. (B-F) Examples of confocal microscopy detection of cyanobacteria in the 20-180 µm size fraction. From top left to bottom right, the displayed channels for each micrograph correspond to cell surface (cyan, AlexaFluor 546 dye), DNA (and the theca in dinoflagellates) (blue, Hoechst dye), cellular membranes (green, DiOC6 dye), chlorophyll autofluorescence (red), bright field, and all merged channels. The size bar at the bottom left of each microscopy image corresponds to 2.5 μm. B) *Trichodesmium* filament. C) *Calothrix* filament outside a chain of the diatom *Chaetoceros* sp. D) *Richelia* filaments inside the diatom *Eucampia cornuta*. E) Picocyanobacterial aggregate. F) Picocyanobacterial symbionts in the dinoflagellate *Ornithocercus thumii*. (G) Correlation analysis between *Trichodesmium* and *Richelia/Calothrix* quantifications by confocal microscopy and *psbO* metagenomic reads in size fraction 20-180 µm. Spearman rho’s correlations coefficients and p-values are indicated. rpkm: reads per kilobase per million mapped reads.

**Figure 7:**
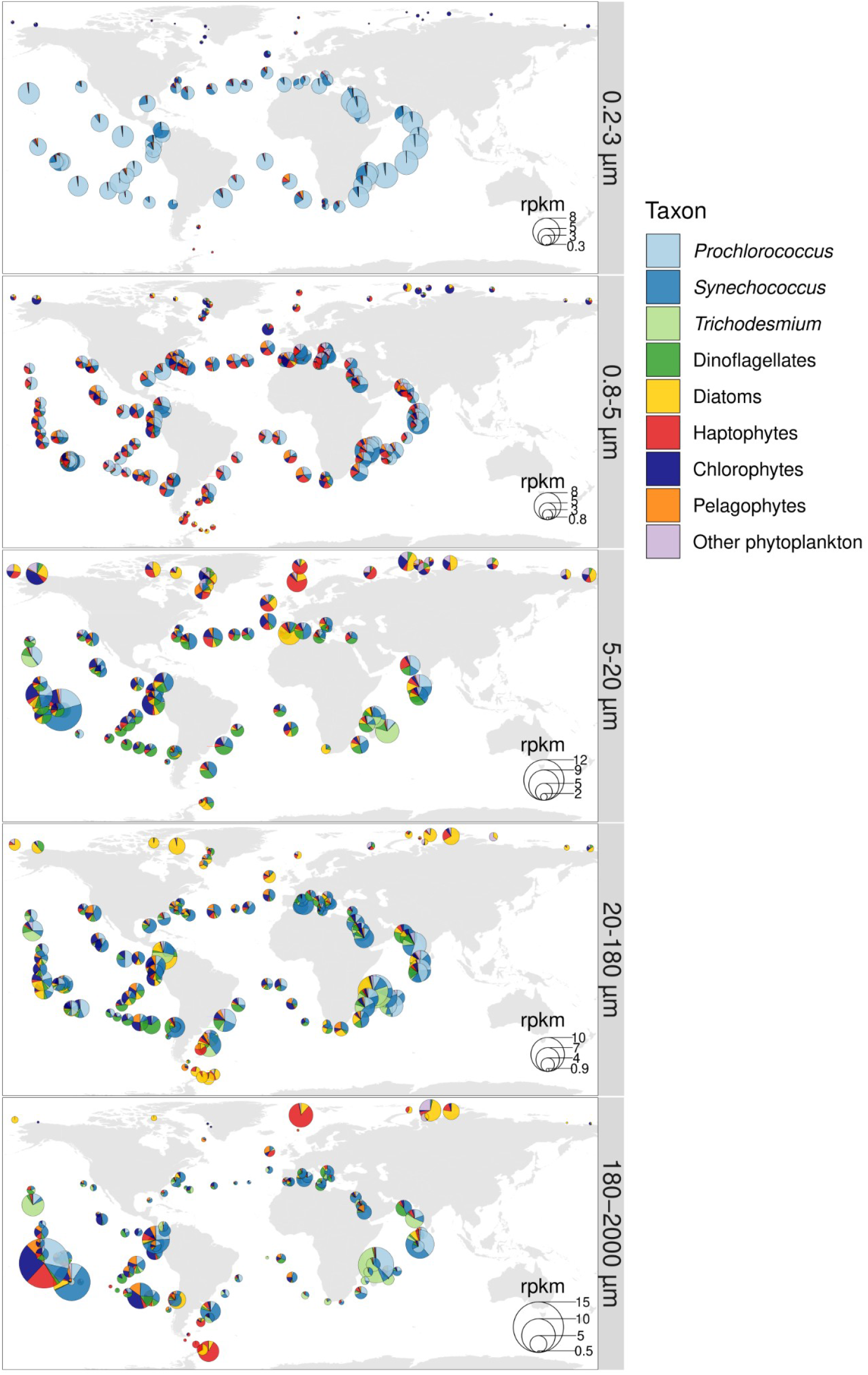
Global biogeographical patterns of marine phytoplankton in surface waters. The pie charts show the *psbO* relative abundance of the main cyanobacteria and eukaryotic phytoplankton in metagenomes derived from different size-fractionated samples. Values are displayed as rpkm (reads per kilobase per million mapped reads). The comparison between the *psbO*-based relative cell abundances versus the patterns corrected by biovolume are displayed in Figure S16. The distribution of the main phytoplankton groups in the size fraction in which they were most prevalent is shown in Figure S17.

To our surprise, we also detected a high abundance of both *Prochlorococcus* and, in particular, *Synechococcus*, in the large size fractions (Figure 6A) across multiple and geographically distinct basins of the tropical and subtropical regions of the world’s ocean (Figure 7). Picocyanobacteria have small cell diameters (<1 μm), and therefore should readily pass through the filters with pore sizes of 5, 20 or 180 μm. Although smaller cells can get caught on larger filters, their abundance should be limited and hence not responsible for the values observed. The reason why a substantial fraction of picocyanobacteria were found in the largest size fractions may be colony formation, symbiosis, attachment to particles, or their grazing by protists, copepods and/or suspension feeders. We examined these possibilities by looking at the *Tara* Oceans confocal microscopy dataset, and found many microscopy images evidencing colony formation and symbiosis in the 20-180 μm size fraction (Figure 6E-F). This is in agreement with the mapping of the *Tara* Oceans metagenomes against a recently sequenced single cell genome of a *Synechococcus* living as a dinoflagellate symbiont (Nakayama et al., 2019). In addition, there are reports of picocyanobacterial symbionts among isolates of planktonic foraminifers, radiolarians, tintinnids, and dinoflagellates (Bird et al., 2017; Foster, Collier, & Carpenter, 2006; Kim, Choi, & Park, 2021; Yuasa, Horiguchi, Mayama, Matsuoka, & Takahashi, 2012) and picocyanobacterial colonies were observed in a regional study based on optical methods (Masquelier & Vaulot, 2008) and in lab cultures (W. Deng, Cruz, & Neuer, 2016; Wei Deng, Monks, & Neuer, 2015).

These results suggest that we should move from the traditional view of *Synechococcus*/*Prochlorococcus* as being exclusively part of picoplankton communities, and instead we should consider them as part of a broader range of the plankton size spectrum (in a similar way as occurs with other small-celled phytoplankton such as the haptophyte *Phaeocystis*; (Beardall et al., 2009; Decelle et al., 2019)). However, it should be borne in mind that these results correspond to estimates of relative cell abundance, and thus the picture is very different when translated to biovolume, due to the large differences in cell size (Figure S16). All in all, our approach allows us to make trans-domain comparisons, which can reveal photosymbiosis and cell aggregates (Figure 6), and allows us to examine the biogeography of the entire phytoplankton community simultaneously (Figures 7, S16 and S17).

## Discussion

We searched for core photosynthetic, single-copy, nuclear genes in genomes and transcriptomes of cultured phytoplankton strains for their use as marker genes. Of the five resulting candidates, *psbO* emerged as the most suitable due to its lack of non-photosynthetic homologs (but note that the other genes could be incorporated in future studies by discarding non-photosynthetic homologs by phylogenetic and/or sequence similarity methods). We applied this new approach by retrieving *psbO* sequences from the metagenomes generated by *Tara* Oceans, and successfully validated it using the optical determinations from the same expedition.

We also quantified the biases of “traditional” molecular approaches as compared to the optical methods. The 16S miTags approach avoids PCR biases and seems to be little affected by copy variability of the 16S gene, the plastid genome and the chloroplast, probably because only the 0.2-3 size fraction was analysed in the current work, where most picoplankton cells only have a single chloroplast. It would be a future scope to analyse larger size fractions where abundant taxa have multiple plastids (e.g., centric diatoms) or divergent 16S genes difficult to align (e.g., dinoflagellates). When compared with 18S metabarcoding data, our approach yields lower abundances for diatoms and dinoflagellates at the expense of higher abundances of haptophytes, chlorophytes and pelagophytes. These results were remarkably consistent with those obtained by microscopy. To disentangle the effect of PCR-bias versus copy number in the patterns of 18S metabarcoding, the next step will be to generate 18S miTags from the analysed metagenomes. It is important to take into account that not all analyses are affected by the biases introduced by traditional molecular methods, as we showed for the Shannon index.

While our work demonstrated that *psbO* reflects the relative cell abundance of phytoplankton, some previous studies suggested that rRNA genes reflect the relative biovolume of the corresponding taxa. However, there is still no clear consensus for rRNA genes as proxies of biovolume. Here, we did not observe major differences between rRNA gene or *psbO* when correlated against optical-based biovolumes.

In addition to correcting for the abovementioned biases, we revealed unexpected ecological features missed by 18S metabarcoding. For example, our trans-domain comparison detected picocyanobacteria in high numbers in large size fractions, which was supported by the observation of numerous images of picocyanobacterial aggregates and endosymbionts in the *Tara* Oceans imaging dataset. Moreover, when the analysis of metagenomes includes housekeeping marker genes (in addition to photosynthetic genes), we observed small dinoflagellates (0.8-5 µm) to be mainly heterotrophic, while those in the larger size communities (>5 µm) to be mainly photosynthetic.

In addition to metagenomes, we also analysed *psbO* in metatranscriptomes, where dinoflagellates stood out from the rest due to their much higher *psbO* abundance ratio of mRNA abundance to gene copy number. It will be of interest to analyse if this reflects higher ‘photosynthetic activity’ or if it is an effect of their predominant post-transcriptional regulation (Cohen et al., 2021; Roy, Jagus, & Morse, 2018).)

The very deep sequencing of the *Tara* Oceans metagenomes (between ∼10^8^ and ∼10^9^ total reads per sample) allowed us to carry out taxonomic analysis based on a unique gene, in spite of dilution of the signal. As reduced DNA sequencing costs are leading to the replacement of amplicon-based methods by metagenome sequencing, we expect the utility of our method to increase in future years. In the short term, a barcode approach using *psbO* primers is a promising cheap alternative, although it will be subject to PCR biases and affected by the presence of introns.

The use of functional genes as taxonomic markers for phytoplankton has been restricted to some surveys (using plastid-encoded genes) (Farrant et al., 2016; Man-Aharonovich et al., 2010a; Paul et al., 2000; Zeidner, Preston, Delong, Massana, Post, Scanlan, & Beja, 2003). This is not the case for other functional groups, such as nitrogen-fixers, which are studied by targeting a gene encoding a subunit of the nitrogenase enzymatic complex (Zehr & Paerl, 1998) and for which extensive reference sequence databases are now available (https://www.jzehrlab.com; (Heller, Tripp, Turk-Kubo, & Zehr, 2014)). To facilitate the incorporation of *psbO* into future molecular-based surveys, we have generated a database of >18,000 annotated *psbO* sequences (https://www.ebi.ac.uk/biostudies/studies/S-BSST659; Figure S1). We hope that the release of this data, and the establishment of *psbO* as a new biomarker for quantifying species abundances, opens new perspectives for molecular-based evaluations of phytoplankton communities.

Based on the current analyses, we recommend the use of *psbO* as a proxy of relative cell abundance of the whole phytoplankton community. However, when focusing on either eukaryotes or prokaryotes, Shannon index is robust enough to be based on rRNA genes. Finally, the use of molecular markers (either *psbO* or rRNA genes) as proxies of relative phytoplankton biovolume is not established.

## Acknowledgments

We would like to thank all colleagues from the *Tara* Oceans consortium as well as the Tara Ocean Foundation for their inspirational vision. We also acknowledge Quentin Carradec for his help with genes encoding ribosomal proteins. This project has received funding from the European Research Council (ERC) under the European Union’s Horizon 2020 research and innovation programme (Diatomic; grant agreement No. 835067). Additional funding is acknowledged from the FFEM - French Facility for Global Environment (Fonds Français pour l’Environnement Mondial), and the French Government “Investissements d’Avenir” Programmes MEMO LIFE (Grant ANR-10-LABX-54), Université de Recherche Paris Sciences et Lettres (PSL) (Grant ANR-1253 11-IDEX-0001-02), France Genomique (ANR-10-INBS-09), and OCEANOMICS (Grant ANR-11-BTBR-0008). JJPK acknowledges postdoctoral funding from the Fonds Français pour l’Environnement Mondial. RGD acknowledges a CNRS Momentum Fellowship, awarded 2019-2021. This article is contribution number *** of *Tara* Oceans.

## Author contributions

JJPK and CB designed the project. JJPK conducted the study and performed the primary data analysis and visualization. JJPK compiled the *psbO* gene reference catalog and EP performed the metagenomic mapping on it. RGD carried out the phylogenetic-based annotation of 16S rRNA gene OTUs. FL, SC and CdV helped with the confocal microscopy dataset, AZ and ES with the optical microscopy and JMG and SGA with the flow cytometry. All authors helped interpret the data. JJPK and CB wrote the paper with substantial input from all authors.

## Data availability statement

All datasets analyzed for this study are of public access as described in Table I. The curated *psbO* database was submitted to the EMBL-EBI repository BioStudies (www.ebi.ac.uk/biostudies) under accession S-BSST659.

---

**Figure S1:**
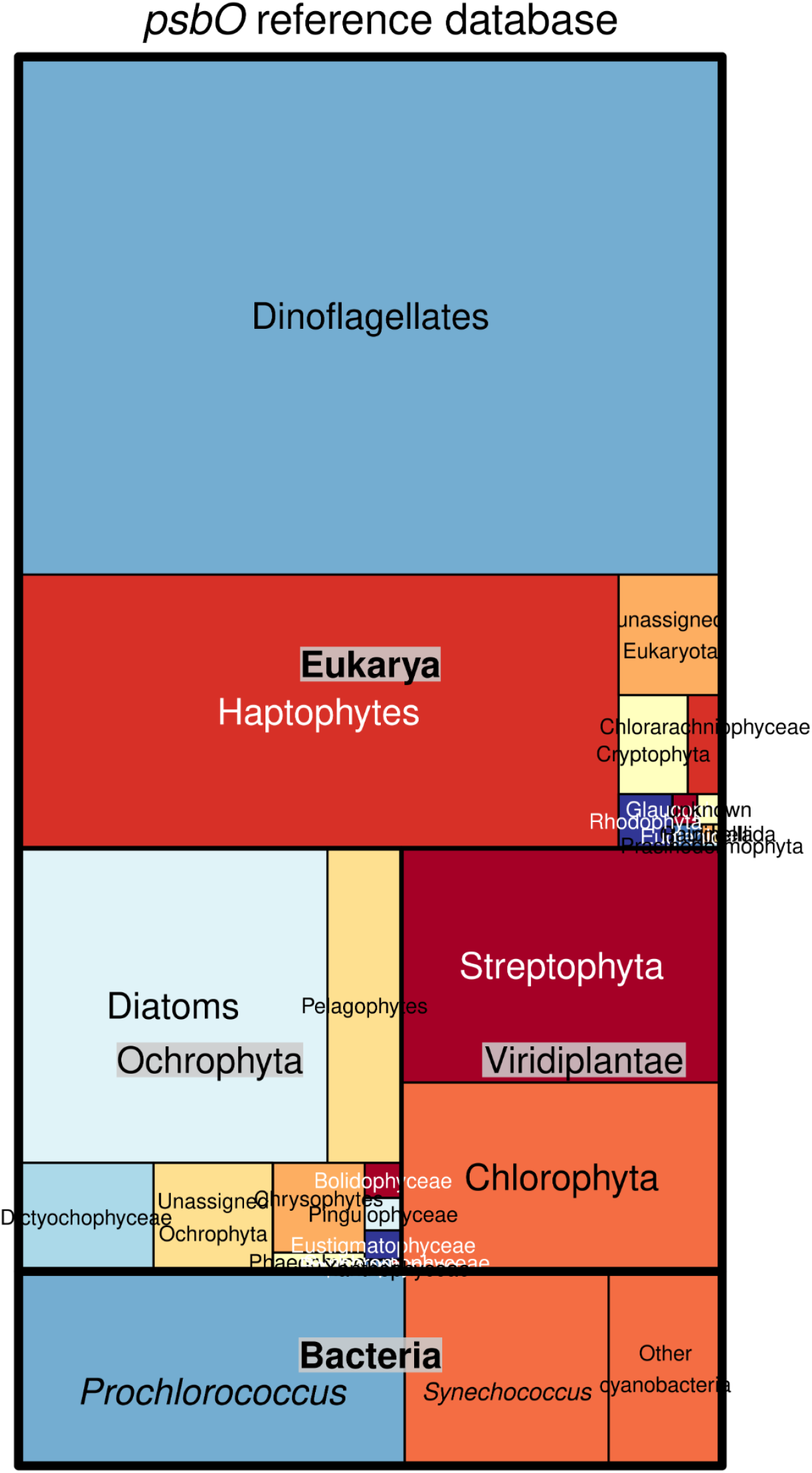
Taxonomic distribution of the curated *psbO* database generated in the current work. It consists of 18,378 unique sequences covering cyanobacteria, photosynthetic protists, macroalgae and land plants. The sequences were retrieved from sequenced genomes and transcriptomes from cultured isolates as well as from the environmental sequence catalogs of Global Ocean Sampling (Rusch et al. 2007) and *Tara* Oceans (Salazar et al. 2019; Carradec et al. 2018; Delmont et al. 2020; 2021). The database can be downloaded from the EMBL-EBI repository BioStudies (www.ebi.ac.uk/biostudies) under accession S-BSST659.

**Figure S2:**
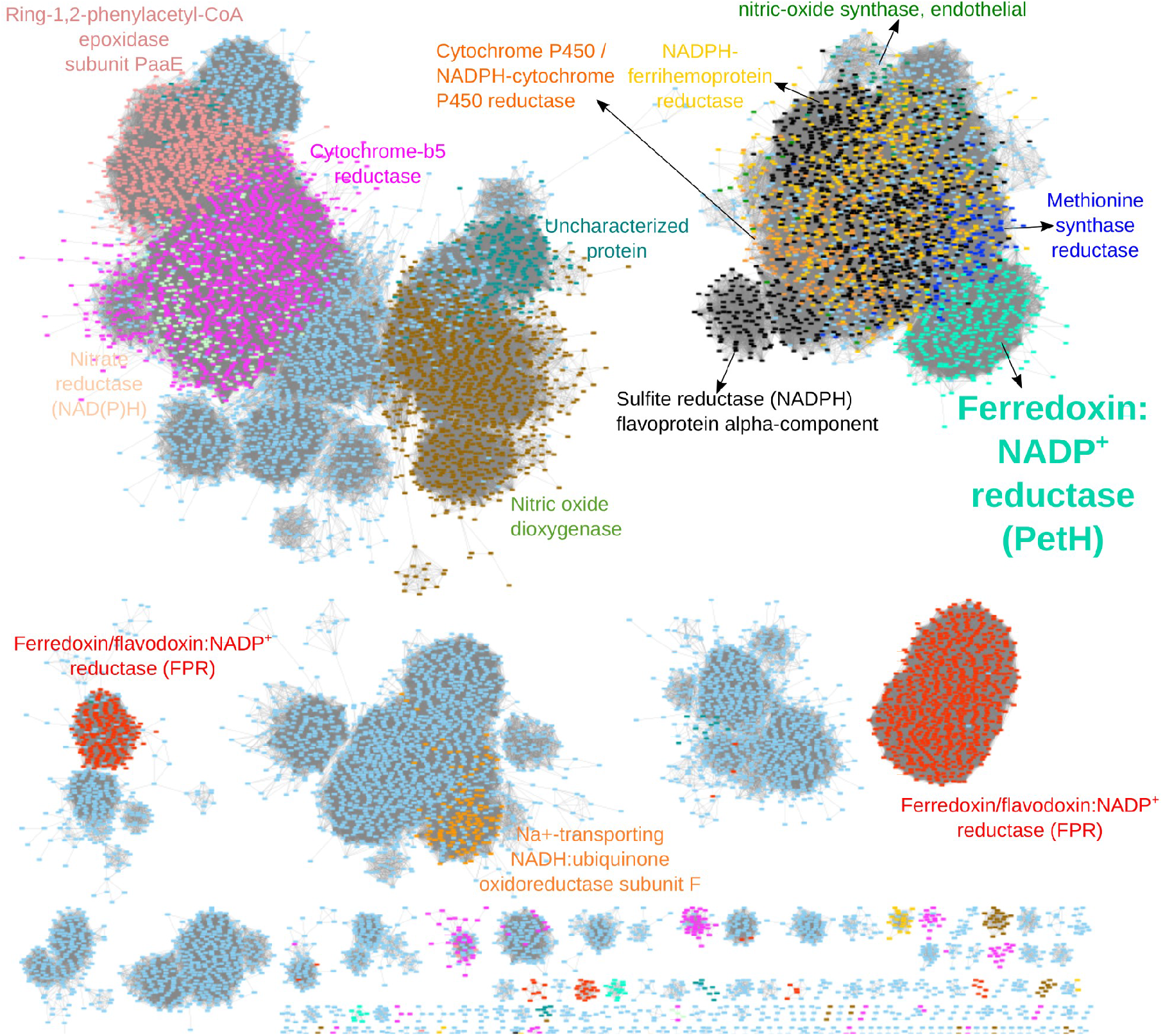
Sequence analysis of ferredoxin:NADP^+^ reductase (PetH) and homologs. Protein similarity network for the Pfam domain NAD_binding_1 (PF00175). Each node corresponds to a representative sequence (clustered at 80% identity by CDHIT) and those sequences with similarity higher than a score cutoff are linked (score cut-off of 22 in blastp alignment). The network was built with sequences retrieved from the literature and from reference genomes and transcriptomes. Nodes are coloured according to their functional assignment based on BlastKOALA. The nodes for FNR are in light green, and includes photosynthetic FNRs as well as FNRs involved in nitrogen metabolism and in non-photosynthetic plastids (Pierella Karlusich & Carrillo 2017 *Phot Res* 134:235–250) and FNR from heterotrophic bacteria acquired by horizontal gene transfer (Catalano Dupuy et al. 2011 *PLoS One* 6:e26736)..

**Figure S3:**
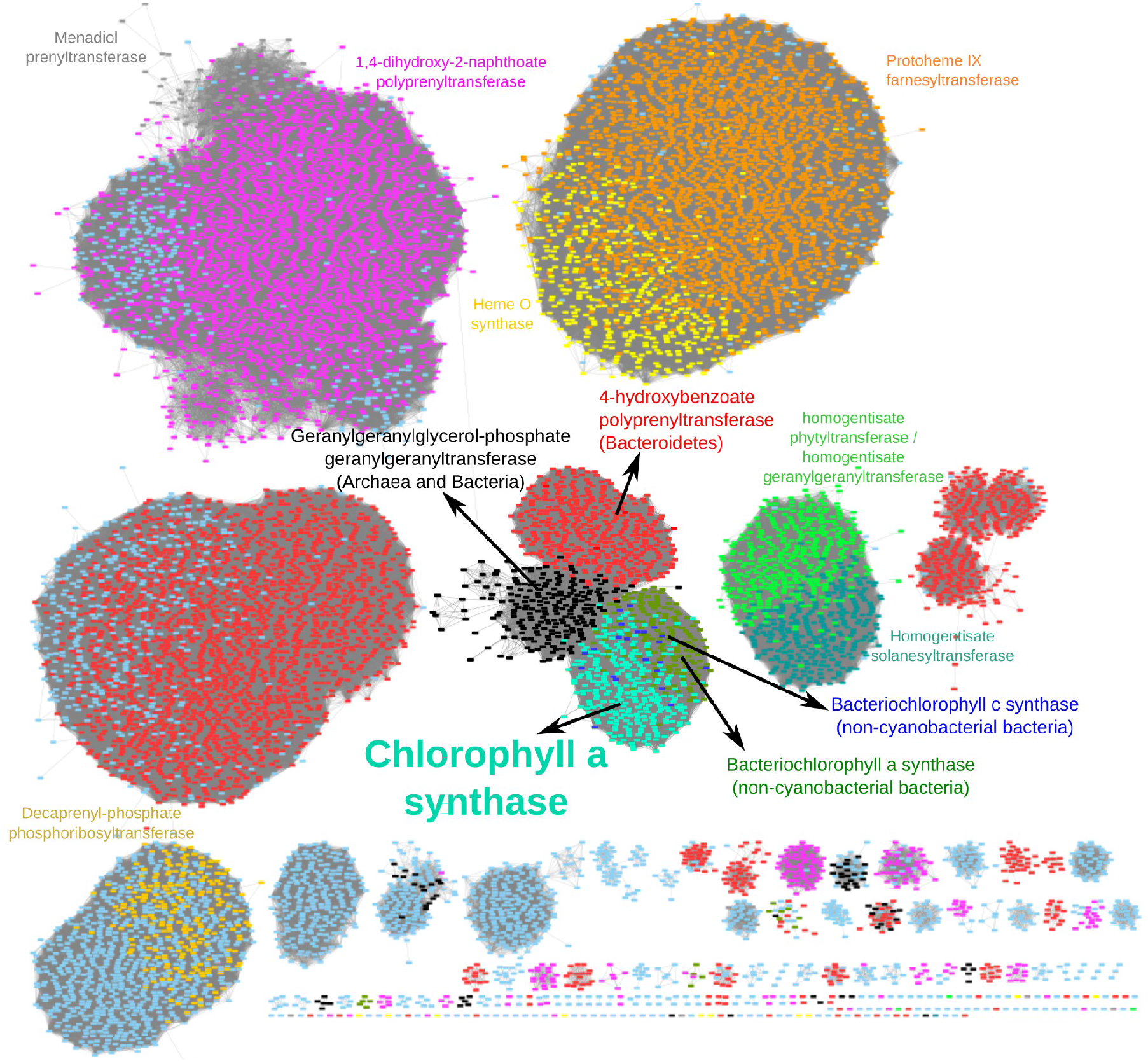
Sequence analysis of chlorophyll *a* synthase (ChlG) and homologs. Protein similarity network for the Pfam domain UbiA (PF01040). Each node corresponds to a representative sequence (clustered at 80% identity by CDHIT) and those sequences with similarity higher than a score cutoff are linked (score cut-off of 25 in blastp alignment). The network was built with sequences retrieved from the literature and from reference genomes and transcriptomes. Nodes are coloured according to their functional assignment based on BlastKOALA.

**Figure S4:**
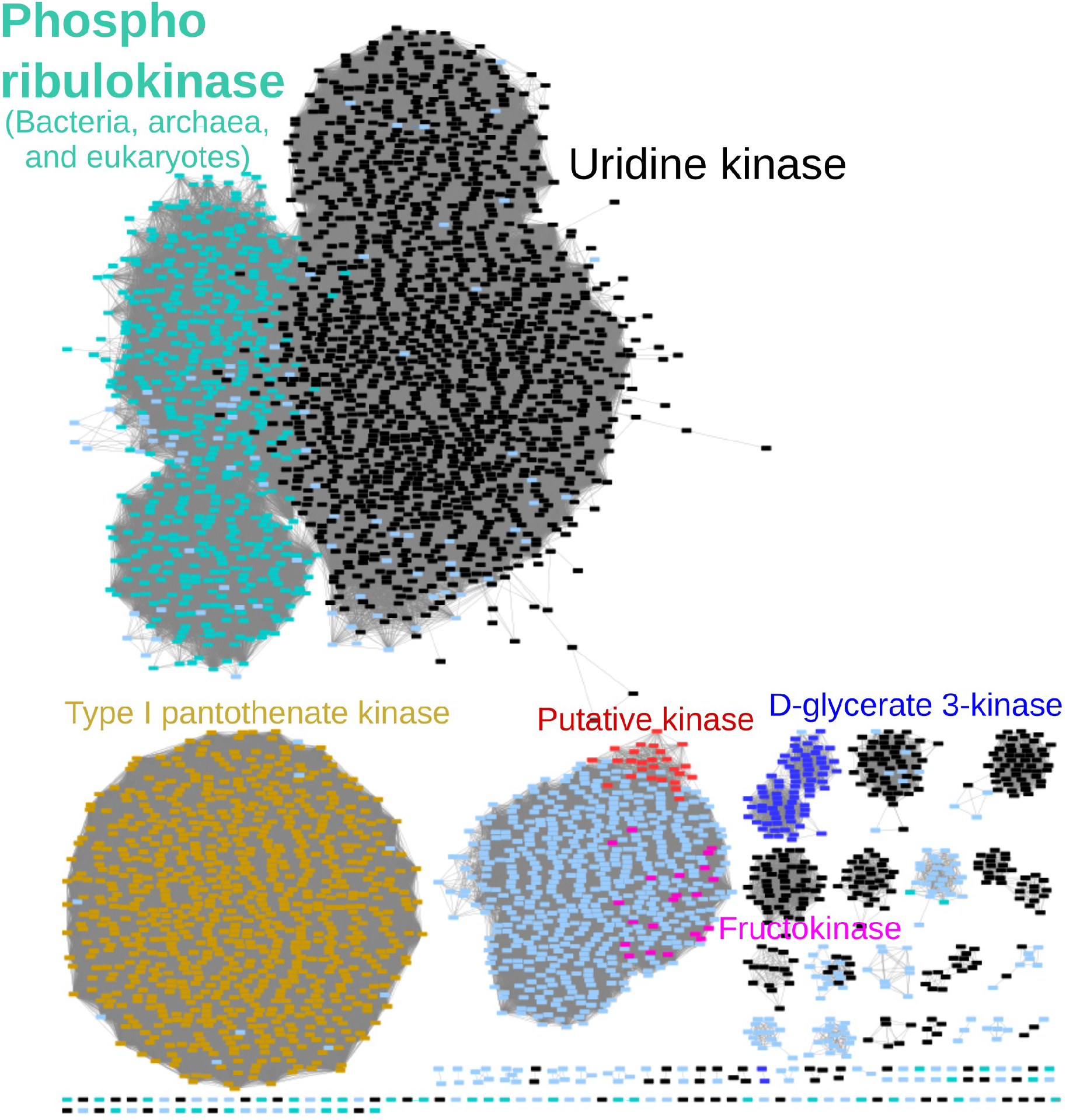
Sequence analysis of phosphoribulokinase (PRK) and homologs. Protein similarity network for the Pfam domain PRK (PF00485). Each node corresponds to a representative sequence (clustered at 80% identity by CDHIT) and those sequences with similarity higher than a score cutoff are linked (score cut-off of 25 in blastp alignment). The network was built with sequences retrieved from the literature and from reference genomes and transcriptomes. Nodes are coloured according to their functional assignment based on BlastKOALA. The nodes for PRK are in light green, and include photosynthetic PRKs as well as those from archaea and non-cyanobacterial bacteria.

**Figure S5:**
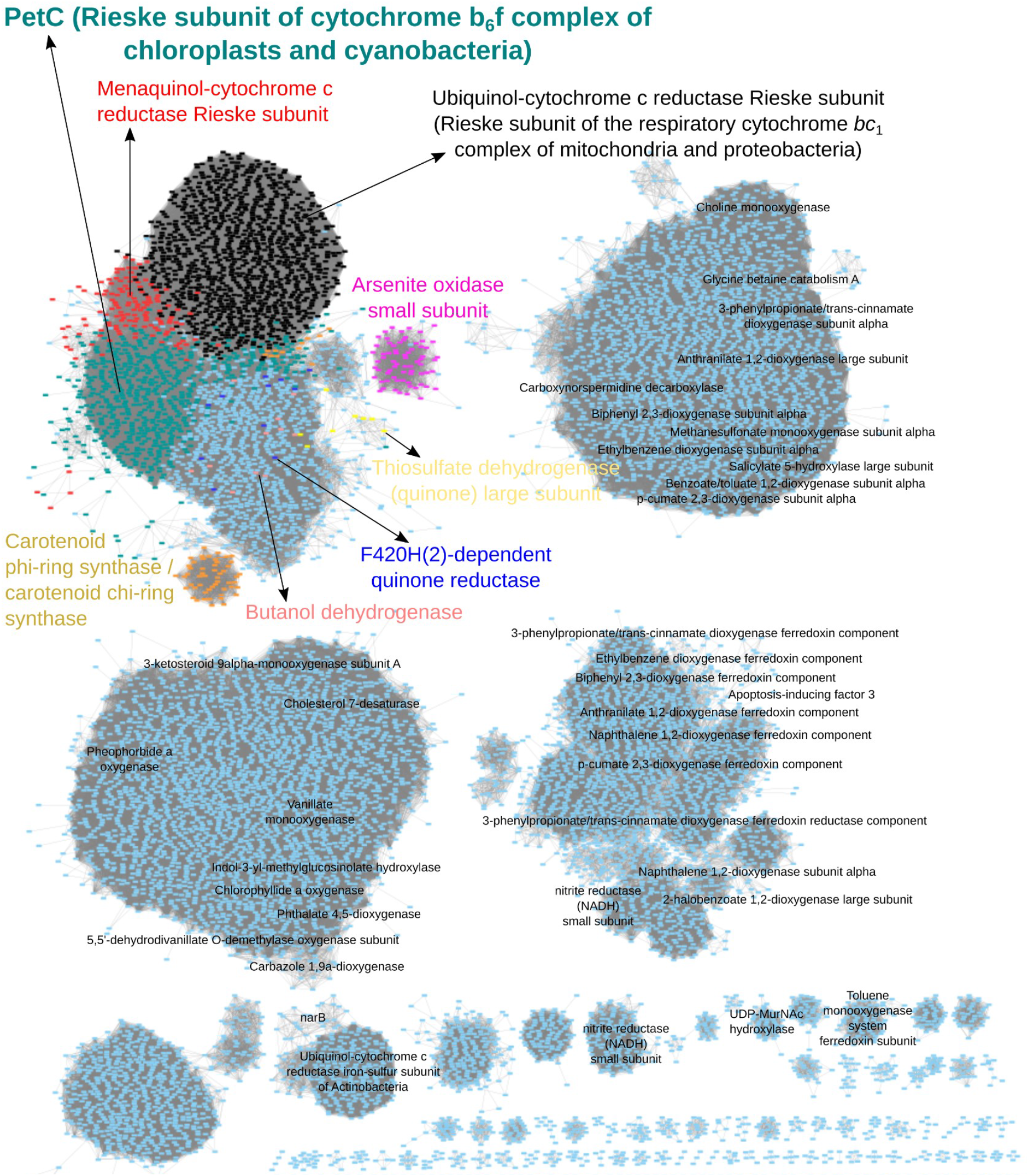
Sequence analysis of PetC (Rieske subunit of the Cytochrome *b*6*f* complex) and homologs. Protein similarity network for the Pfam domain Rieske (PF00355). Each node corresponds to a representative sequence (clustered at 80% identity by CDHIT) and those sequences with similarity higher than a score cutoff are linked (score cut-off of 18 in blastp alignment). The network was built with sequences retrieved from the literature and from reference genomes and transcriptomes. Labels correspond to the functional assignment based on BlastKOALA, and for the cluster of interest nodes are coloured according to the functional assignment of their sequences. The nodes for PetC are in green and those for the Rieske subunit of the respiratory Cytochrome *bc*1 complex are in black.

**Figure S6:**
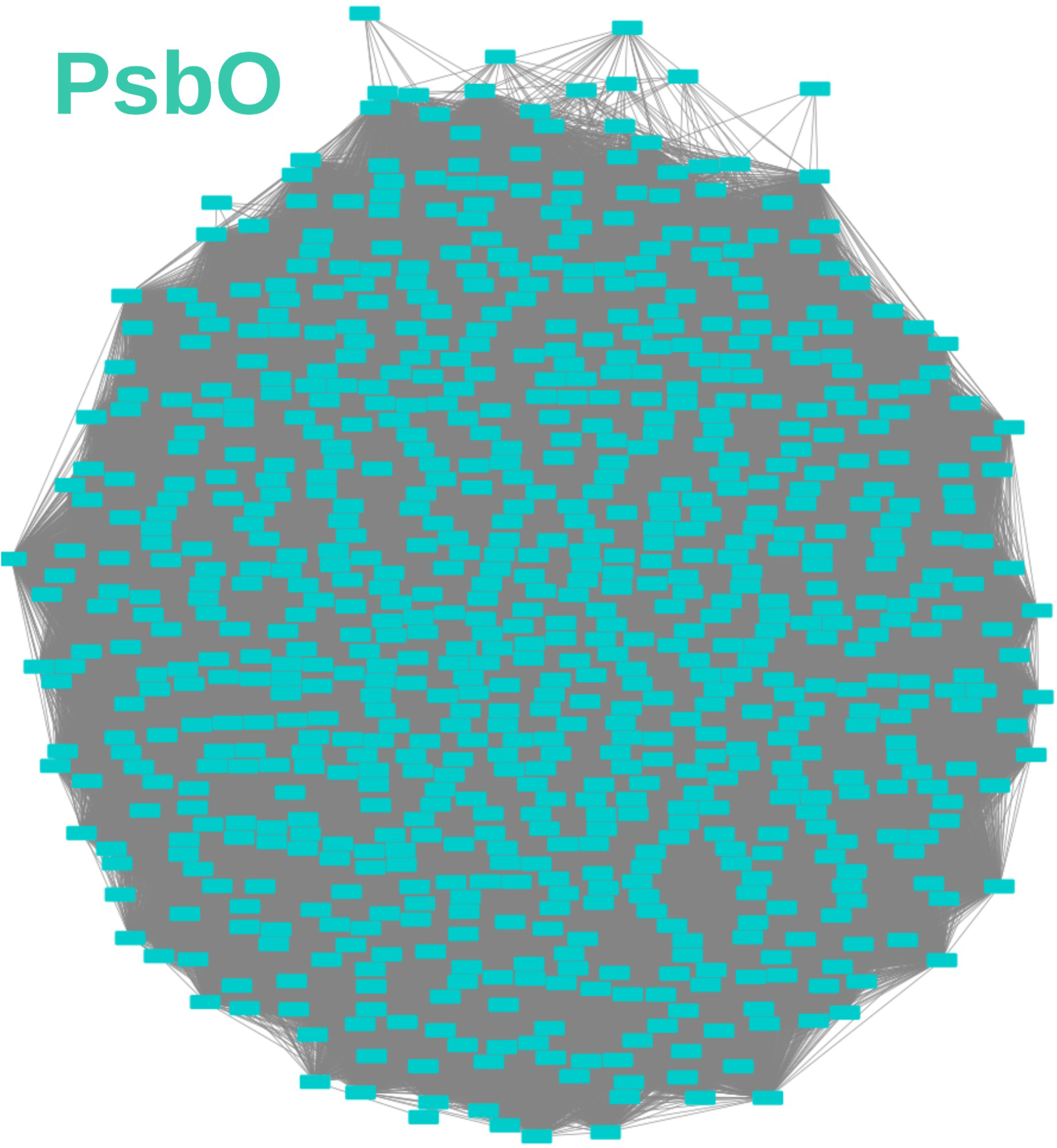
Sequence analysis of PsbO protein. Protein similarity network for the Pfam domain MSP (PF01716). Each node corresponds to a representative sequence (clustered at 80% identity by CDHIT) and those sequences with similarity higher than a score cutoff are linked (score cut-off of 30 in blastp alignment). The network was built with sequences retrieved from the literature and from reference genomes and transcriptomes. Nodes are coloured according to their functional assignment based on BlastKOALA.

**Figure S7:**
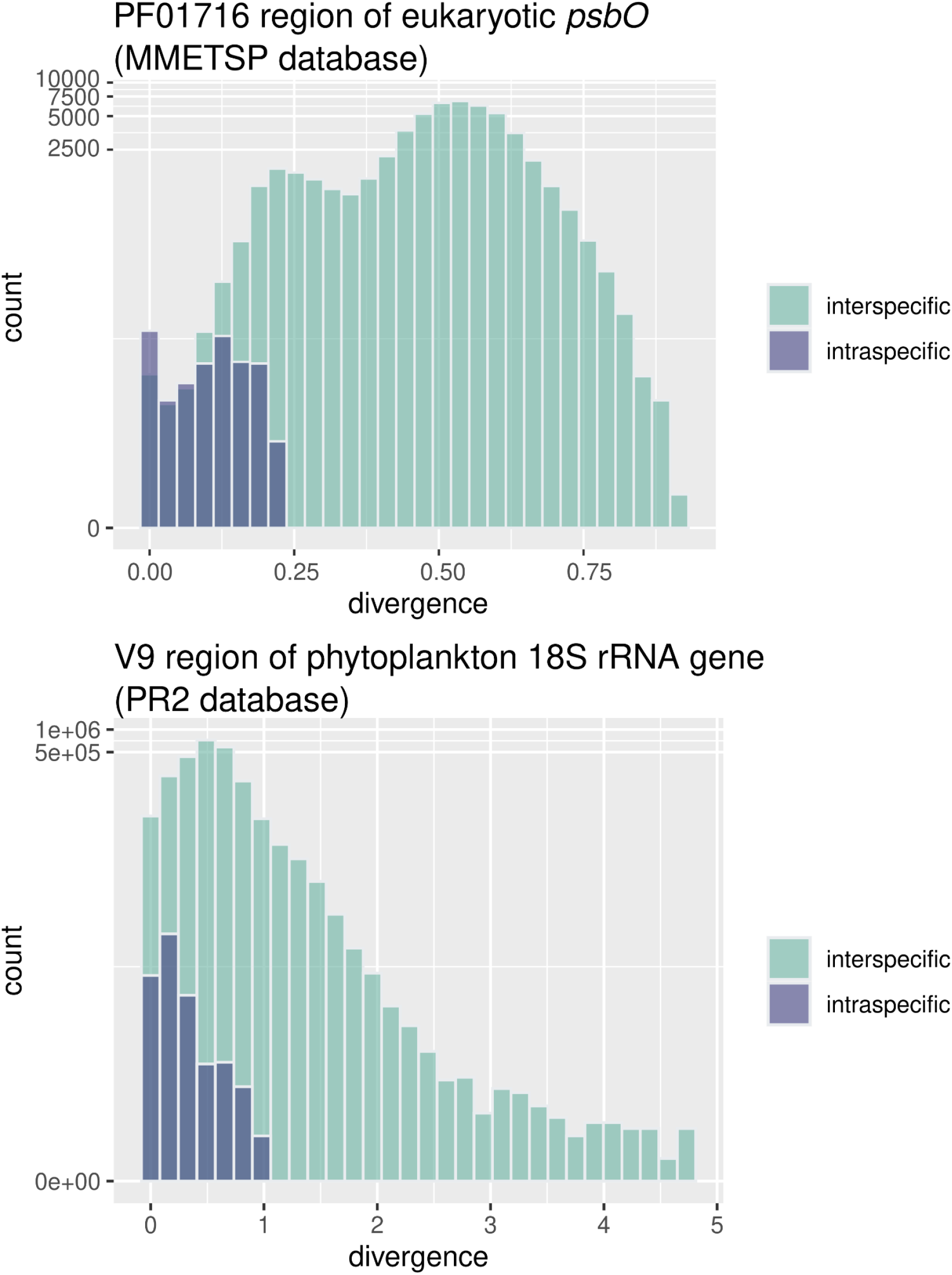
Intraspecific (blue) and interspecific (green) variation in genetic distances of eukaryotic phytoplankton sequences for the region of *psbO* coding for the Pfam domain PF01716 (upper panel) and the V9 region of 18S rRNA gene (lower panel). Sequences were retrieved from MMETSP project (Keeling et al. 2014) for *psbO* and from PR2 database (Guillou et al. 2012) for V9-18S maker. In this later case, the sequence assigned to phytoplankton were selected based on a public functional database available at https://zenodo.org/record/3768951#.YM4odnUzbuE.

**Figure S8:**
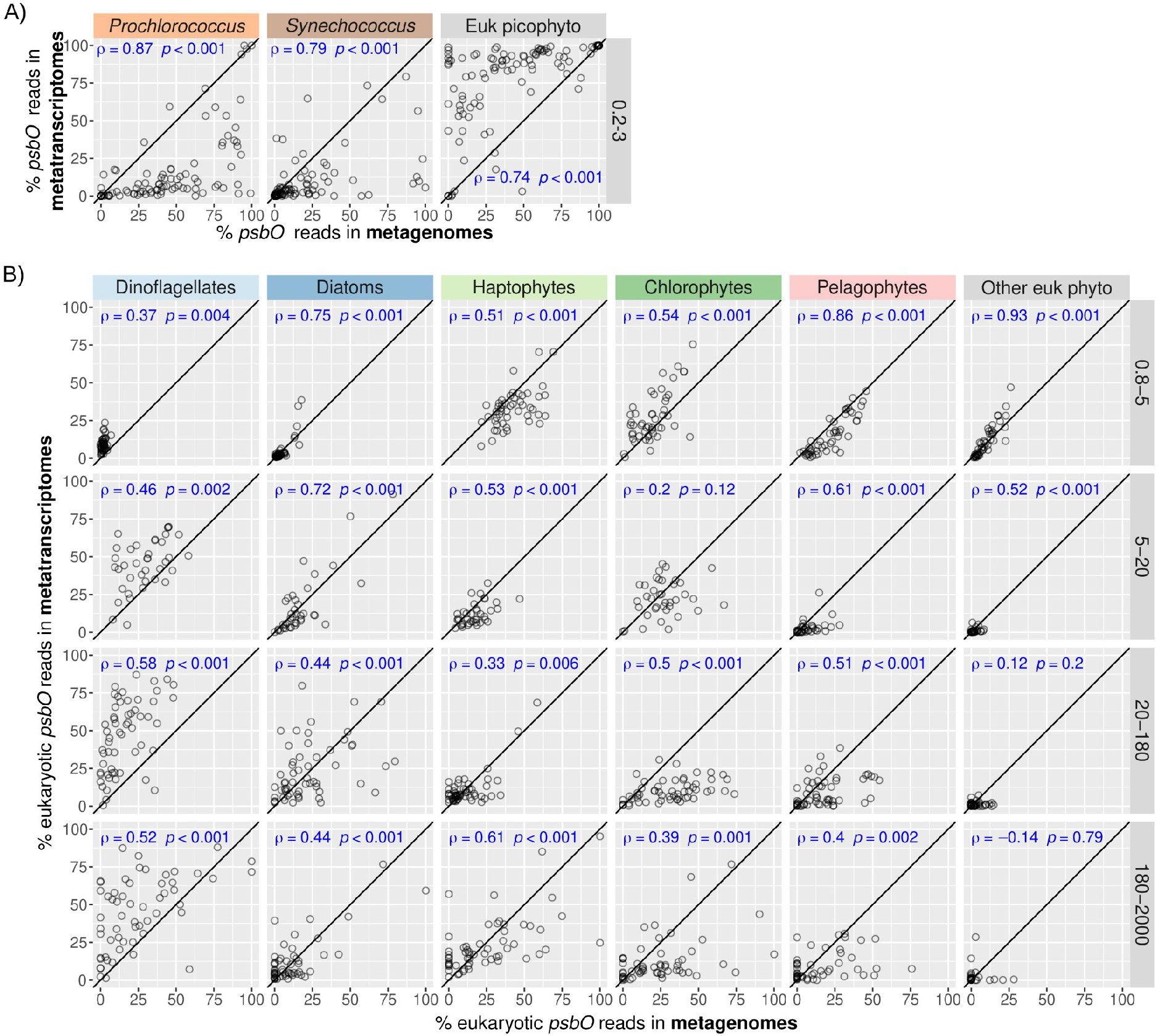
Comparison of relative abundances of *psbO* reads between metagenomes and metatranscriptomes of size fractionated samples. A) Picocyanobacteria and eukaryotic picophytoplankton (size fraction 0.2-3 μm). B) Eukaryotic phytoplankton in the large size fractions (0.8-5 μm, 5-20 µm, 20-180 µm, 180-2000 µm), with metagenomes compared to metatranscriptomes derived from poly-A RNA. Axis are in the same scale and the diagonal line corresponds to a 1:1 slope. Spearman’s rho correlation coefficients and p-values are displayed in blue.

**Figure S9:**
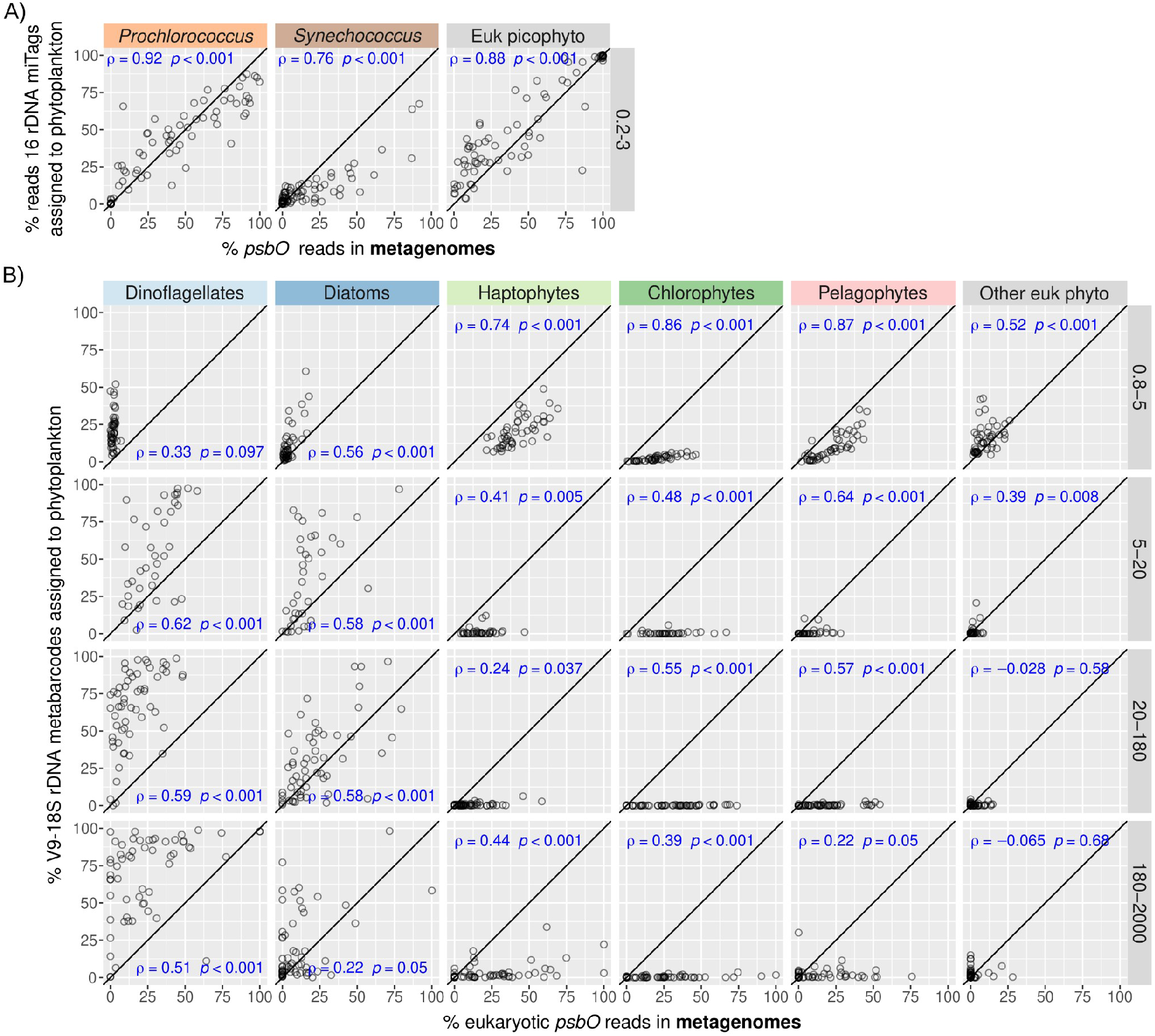
Comparison of relative read abundances of *psbO* and rRNA genes of size fractionated samples. A) Picocyanobacteria and eukaryotic picophytoplankton (0.2-3 μm) were analysed using the relative abundances for 16S rRNA gene miTags and for *psbO* metagenomic reads. B) Eukaryotic phytoplankton were analysed in the large size fractions (0.8-5 μm, 5-20 µm, 20-180 µm, 180-2000 µm) using the relative abundances for V9-18S rRNA gene amplicons and *psbO* metagenomic reads. Axis are in the same scale and the diagonal line corresponds to a 1:1 slope. Spearman’s rho correlation coefficients and p-values are displayed in blue.

**Figure S10:**
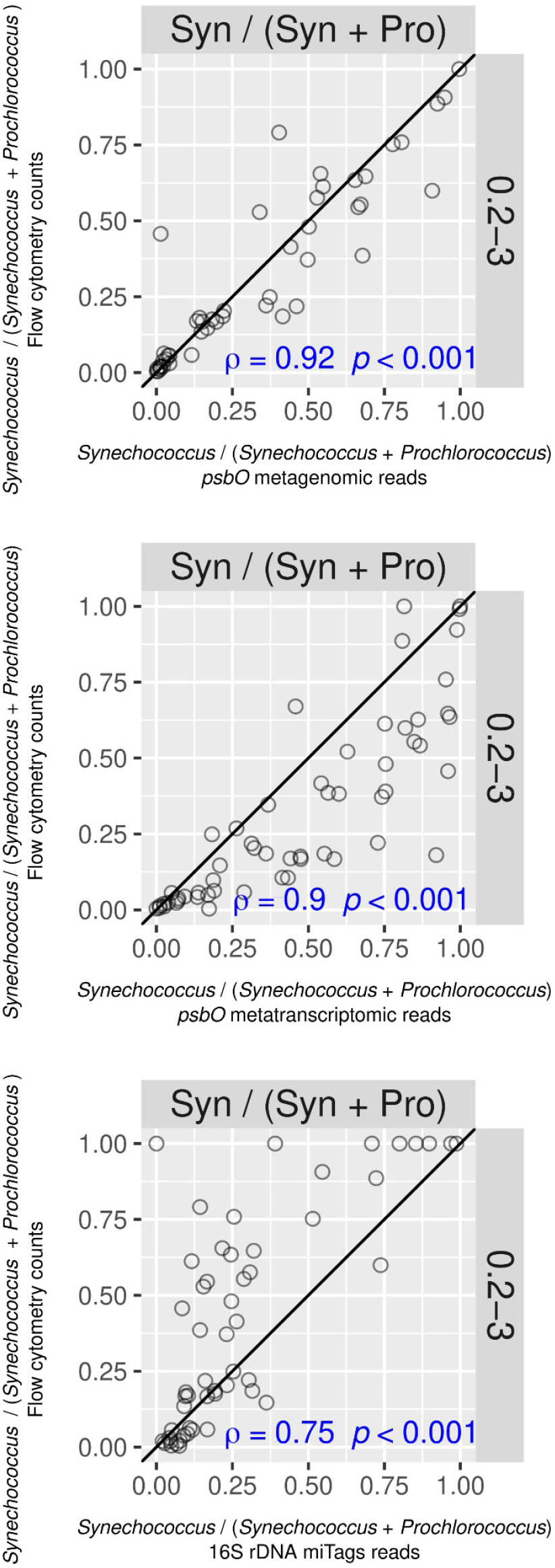
Correlation between the abundance ratio of the picocyanobacteria *Synechococcus* and *Prochlorococcus* obtained with different methodologies. The vertical axis corresponds to the ratio based on flow cytometry while the horizontal axis corresponds to the ratio based on *psbO* metagenomic reads (upper plot) or *psbO* metatranscriptomic reads (middle plot) or 16S miTAGs reads (bottom plot). Axis are in the same scale and the diagonal line corresponds to a 1:1 slope. Spearman’s rho correlation coefficients and p-values are displayed in blue.

**Figure S11:**
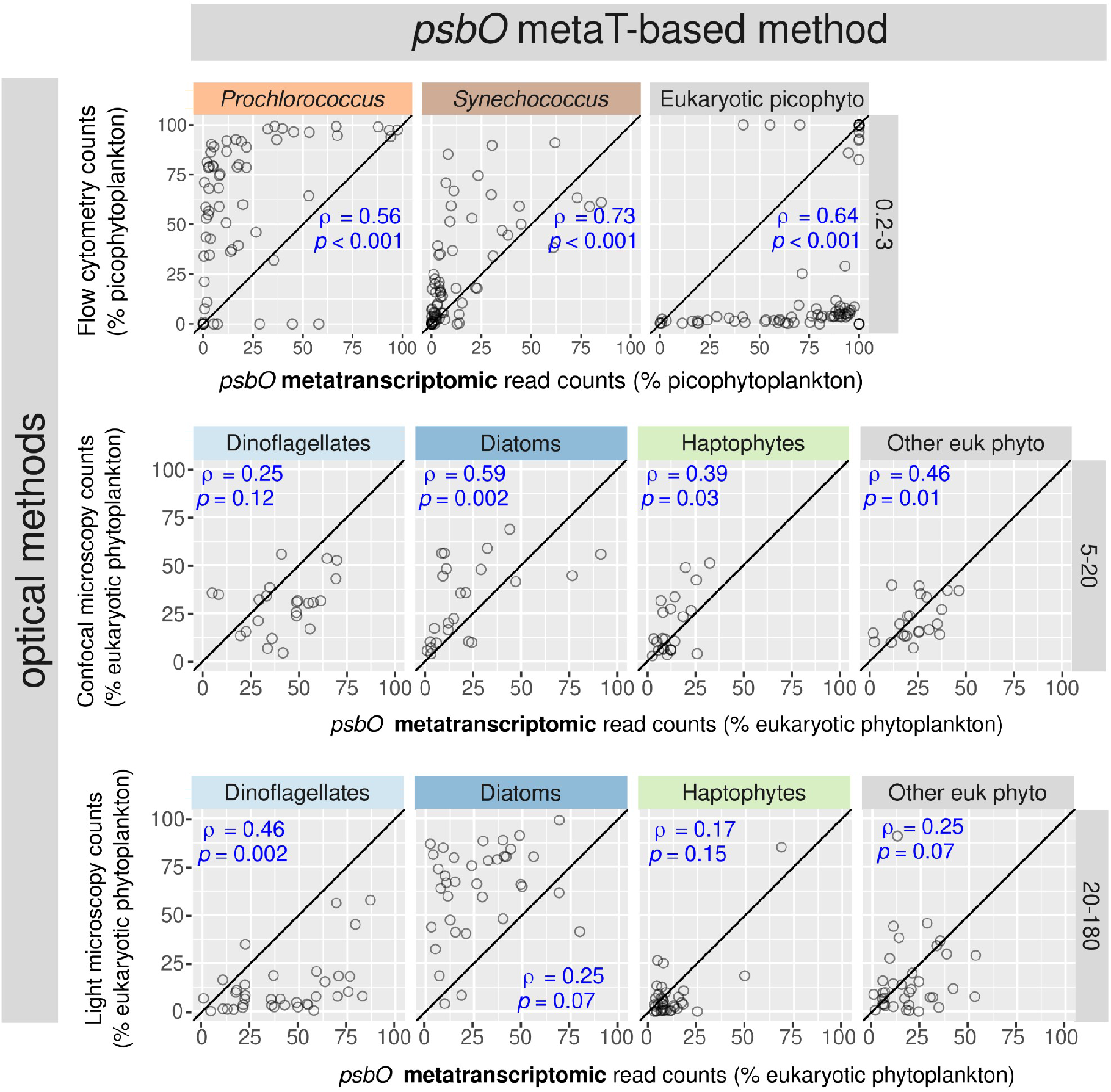
Correlation between relative abundances based on *psbO* metatranscriptomic reads against those based on optical quantifications of different phytoplankton groups. Upper panel: metatranscriptomic *psbO* relative abundances from picophytoplankton (size fraction 0.2-3 µm) were compared with flow cytometry counts (values displayed as % total abundance of picophytoplankton). Middle and lower panels: metatranscriptomic *psbO* relative abundances of eukaryotic phytoplankton were compared with confocal microscopy counts from size fraction 5-20 μm and light microscopy counts from size fraction 20-180 µm (values displayed as % total abundance of eukaryotic phytoplankton). Spearman correlation coefficients and p-values are displayed in blue. Axis are in the same scale and the diagonal line corresponds to a 1:1 slope.

**Figure S12:**
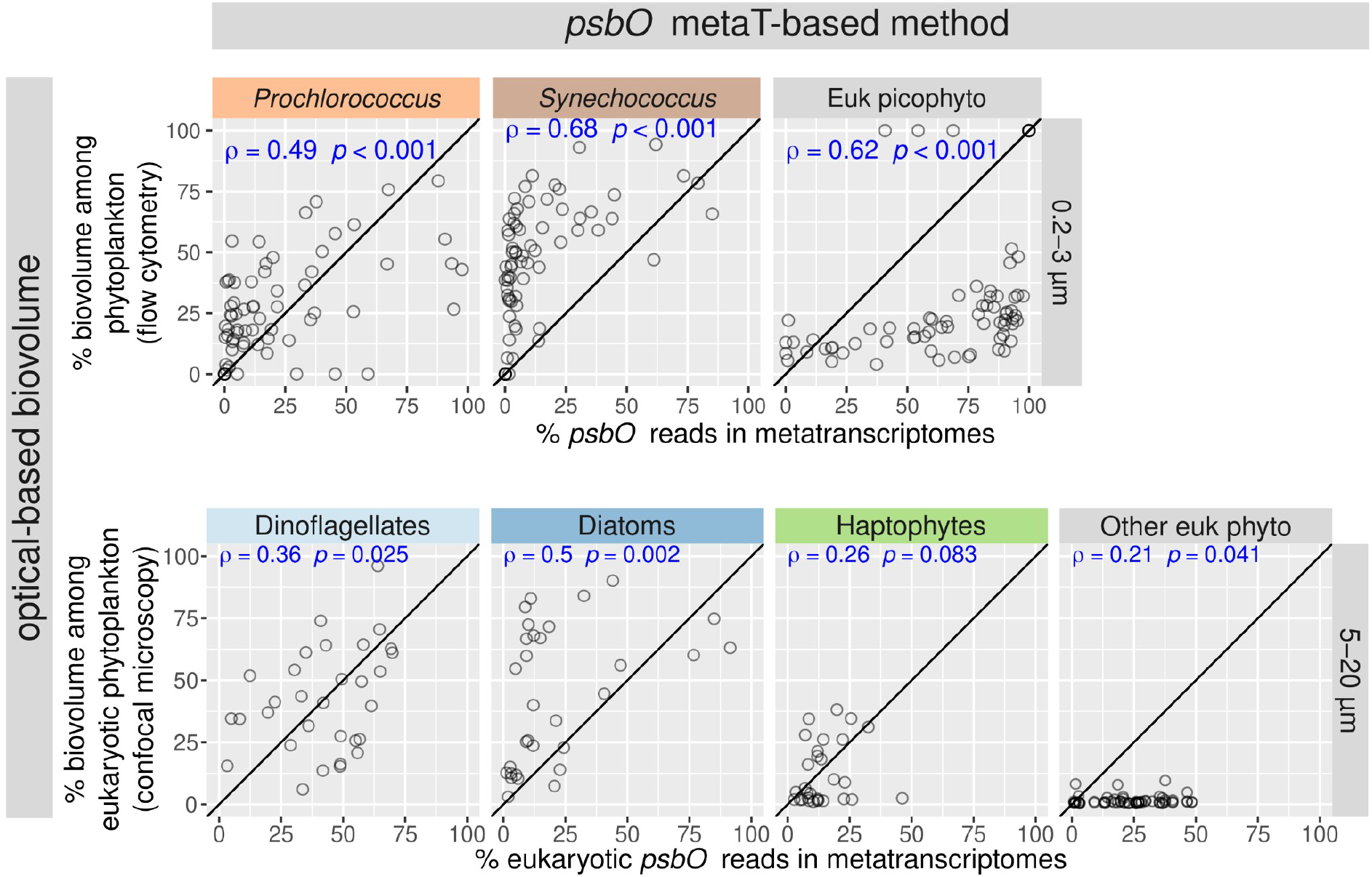
Correlation between relative biovolume (based on optical methods) and relative abundances based on *psbO* metatranscriptomic reads. The upper panel shows the correlations for picophytoplankton (size fraction 0.2-3 µm). The vertical axis corresponds to the relative biovolume based on flow cytometry (values displayed as % total biovolume of picophytoplankton), while the horizontal axis corresponds to relative read abundance based on *psbO* metatranscriptomic reads. The lower panel shows the correlations for nanophytoplankton (size fraction 5-20 μm). The vertical axis corresponds to the relative biovolume based on confocal microscopy quantification (values displayed as % total abundance of eukaryotic phytoplankton), while the horizontal axis corresponds to relative read abundance based on eukaryotic *psbO* metatranscriptomic reads. Spearman correlation coefficients and p-values are displayed in blue. Axis are in the same scale and the diagonal line corresponds to a 1:1 slope.

**Figure S13:**
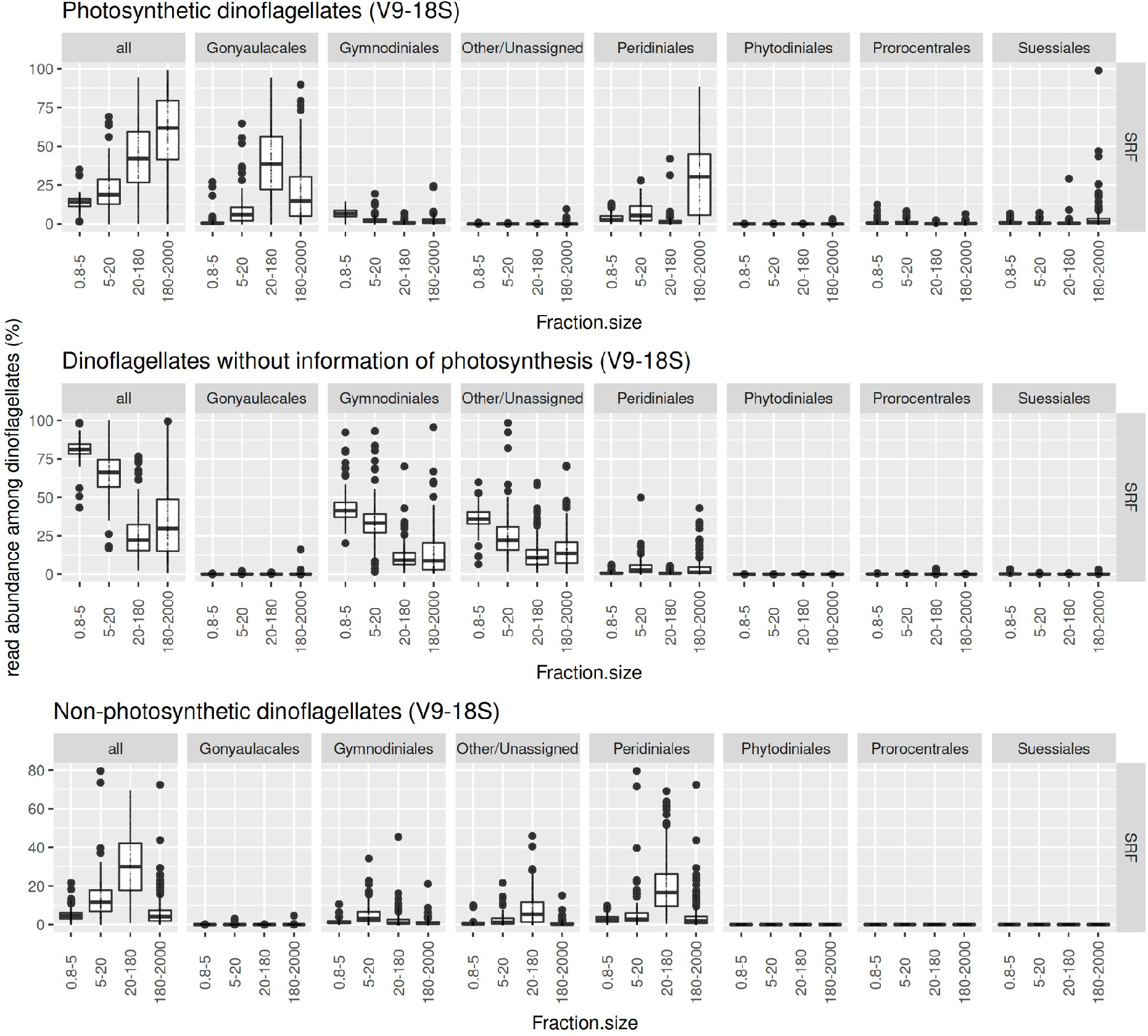
Variations in the abundance of phototrophs *vs* heterotrophs in the dinoflagellate community of different size fractions based on V9-18S rRNA gene metabarcoding. The plots show the relative abundance of dinoflagellates containing chloroplasts (upper panel) or not (lower panel) as well as those that cannot be classified (middle panel). Note that most of the reads that cannot be reliably classified as a chloroplast-containing taxon correspond to those reads mapping OTUs assigned either as “unknown dinoflagellate” or Gymnodiales order. A description of the trait classification can be found at http://taraoceans.sb-roscoff.fr/EukDiv/ and the trait reference database is available at https://zenodo.org/record/3768951#.YM4odnUzbuE.

**Figure S14:**
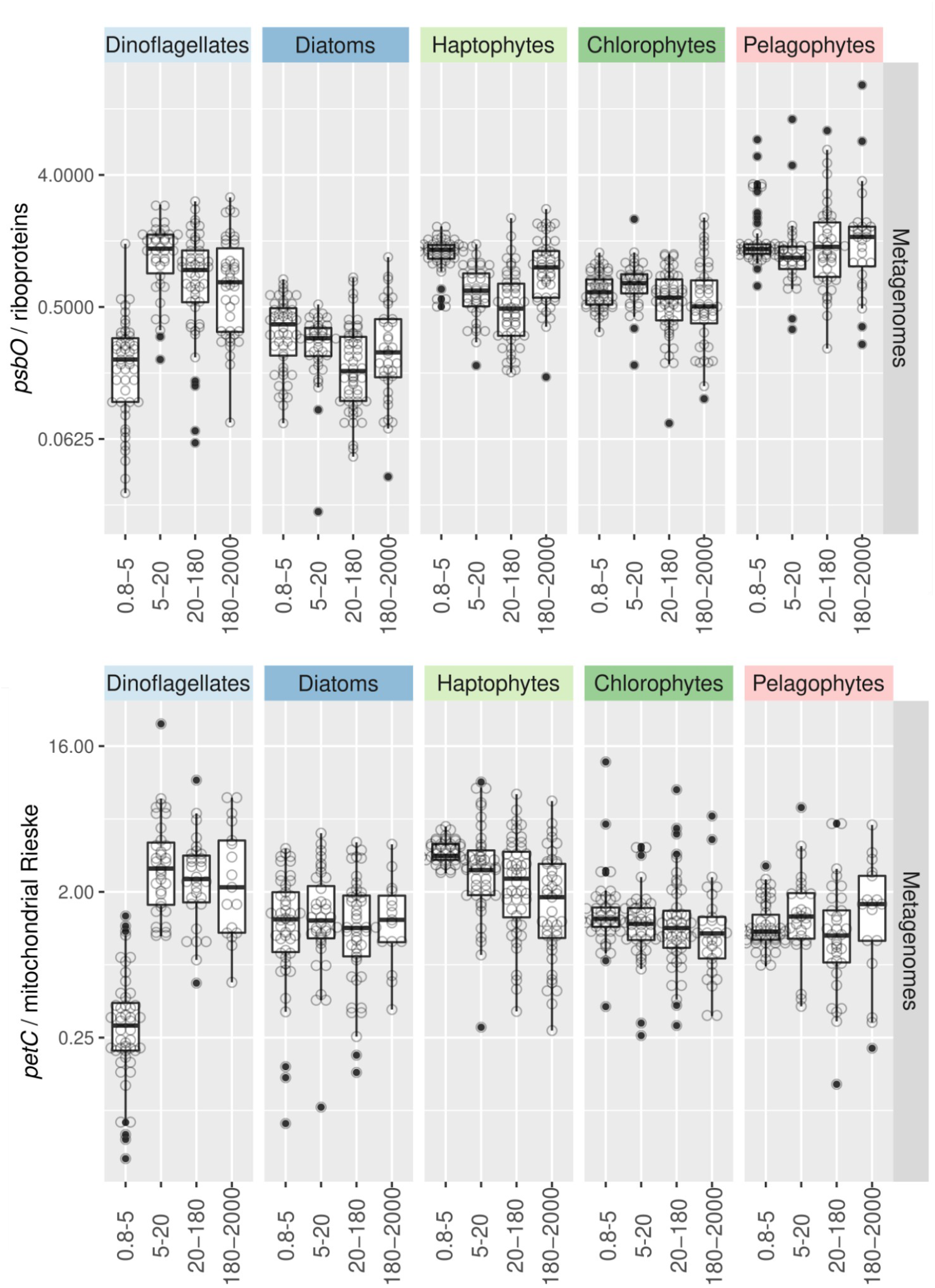
Variations in the abundance of phototrophs *vs* heterotrophs across size fractions based on different marker genes in metagenomes. The estimation were based on the ratio of metagenomic reads of photosynthetic *vs* housekeeping single-copy nuclear-encoded genes: *psbO* vs genes coding for ribosomal protein (upper panel), and the genes coding for the Rieske subunits of the Cyt *bc*-type complexes from chloroplasts and mitochondria (i.e., *petC* and its mitochondrial homologue) (lower panel).

**Figure S15:**
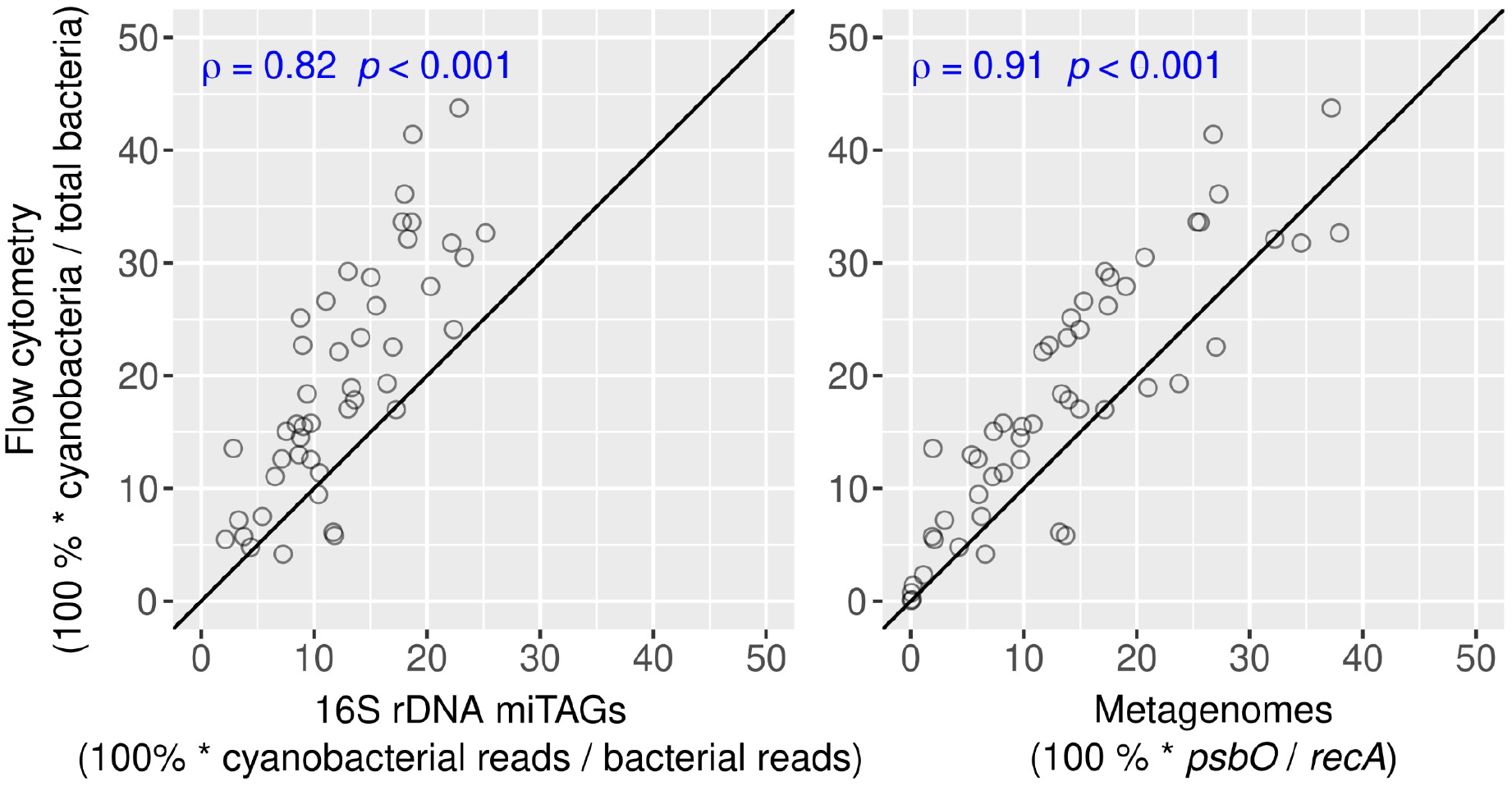
Congruence in the relative abundance of phototrophs among bacterioplankton in 0.2-3 size fraction based on different methods. The vertical axis corresponds to the relative cell abundance based on flow cytometry while the horizontal axis corresponds to the relative read abundance based on 16S miTags (left) or on the ratio of *psbO* to *recA* in metagenomes (right). Spearman correlation coefficients and p-values are displayed in blue. Axis are in the same scale and the diagonal line corresponds to a 1:1 slope.

**Figure S16:**
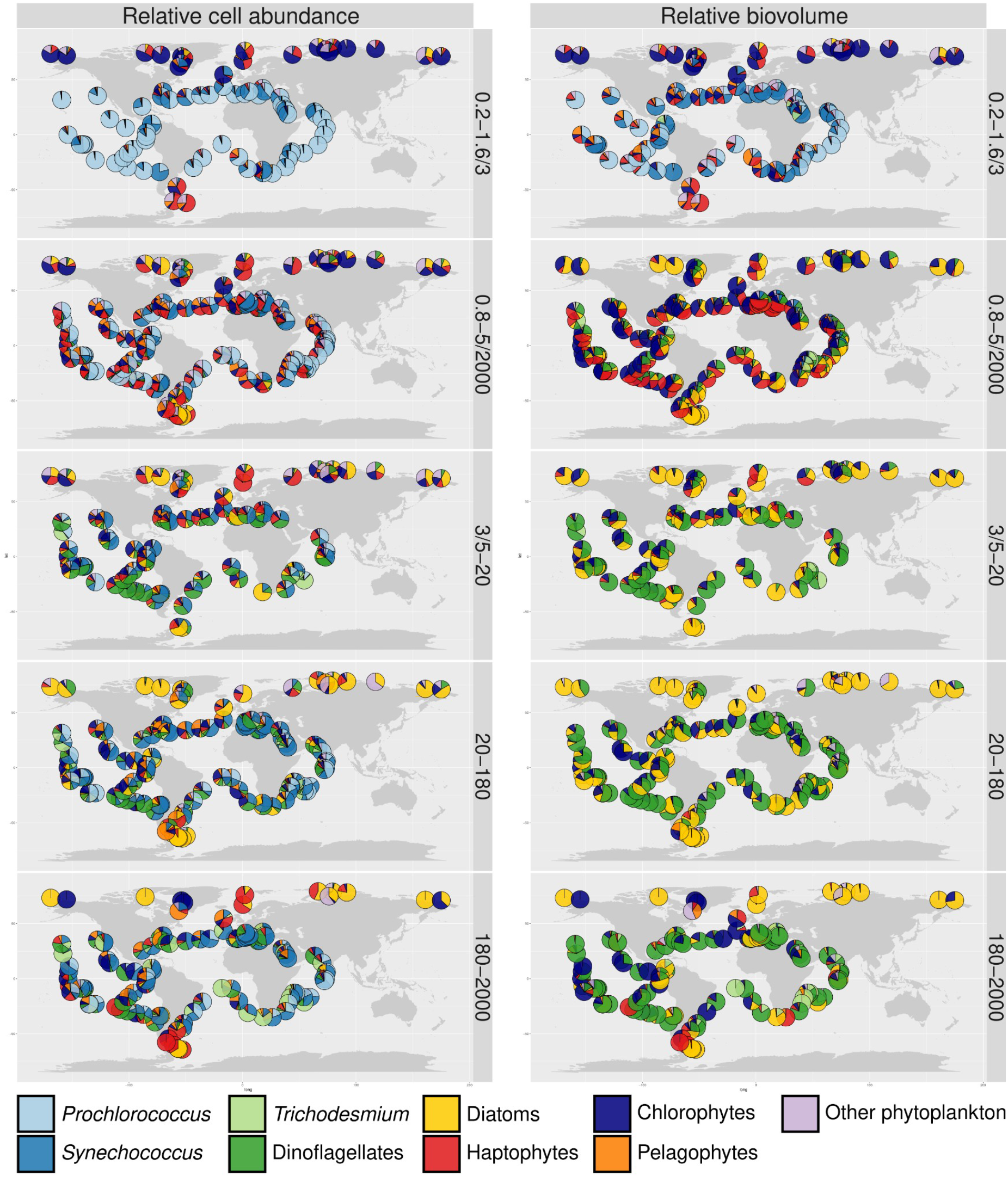
Global biogeographical patterns of relative cell abundance and biovolume for marine phytoplankton in surface waters. A) Relative cell abundances of the main cyanobacteria and eukaryotic phytoplankton based on *psbO* counts in metagenomes derived from different size-fractionated samples. B) Relative biovolume of the main cyanobacteria and eukaryotic phytoplankton based on *psbO* counts correctected by the mean cell biovolume for each taxon (based on optical measurements in *Tara* Oceans samples).

**Figure S17:**
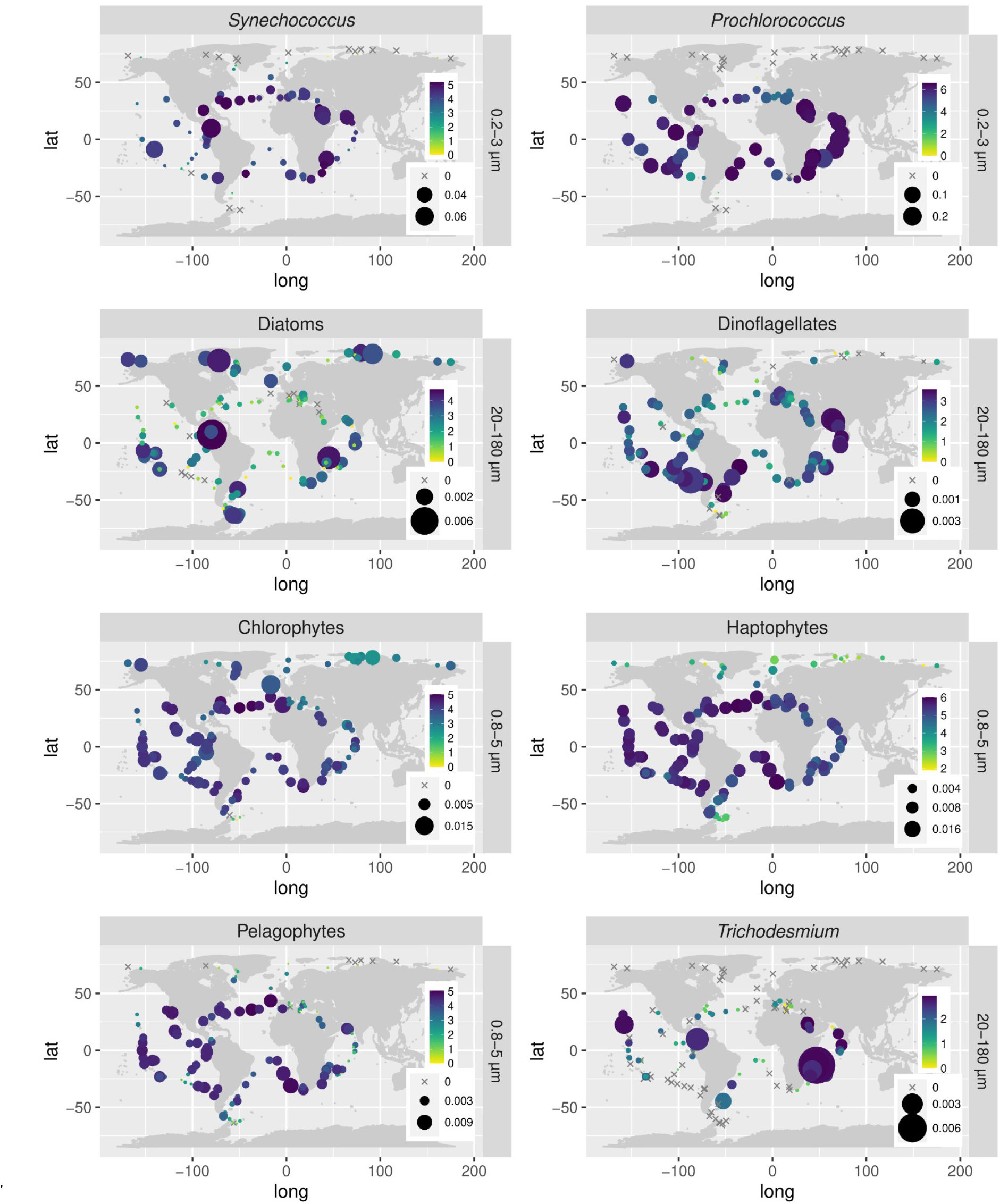
Global biogeographical patterns for main groups of marine phytoplankton in surface waters. The bubbles sizes vary according to the *psbO* relative abundance of the main cyanobacteria and eukaryotic phytoplankton in metagenomes, while color code corresponds to the Shannon index values. Relative abundance values are displayed as rpkm (reads per kilobase per million mapped reads). Only the size fraction where the corresponding taxon was prevalent is shown: 0.2-3 μm for picocyanobacteria, 20-180 μm for diatoms and dinoflagellates, and 0.8-5 μm for chlorophytes, haptophytes and pelagophytes. The corresponding analysis for the whole phytoplankton community in each size fraction is displayed in Figure 7.

